# Influence of claustrum on cortex varies by area, layer, and cell type

**DOI:** 10.1101/2022.02.22.481532

**Authors:** Ethan G. McBride, Saurabh R. Gandhi, Jacqulyn R. Kuyat, Douglas R. Ollerenshaw, Anton Arkhipov, Christof Koch, Shawn R. Olsen

## Abstract

The claustrum is a small subcortical structure with widespread connections with disparate regions of the cortex. These far-reaching projections have led to many hypotheses concerning its function. However, we know little about how claustrum input affects neural activity in cortex, particularly beyond frontal areas. Here, using optogenetics and multi-regional Neuropixels recordings from over 15,000 neurons in awake mice, we demonstrate that the effect of claustrum input differs depending on brain area, layer, and cell type. Brief claustrum stimulation produces approximately 1 spike per claustrum neuron, which affects many fast-spiking (FS; putative inhibitory) but very few regular-spiking (RS; putative excitatory) cortical neurons. Prolonged claustrum stimulation affects many more cortical FS and RS neurons. More inhibition occurs in frontal regions and deeper layers, while more excitation occurs in posterior regions and superficial layers. These differences imply that the function of claustrum input to cortex depends on the area, supporting the idea that claustro-cortical circuits are organized into functional modules.

## INTRODUCTION

The claustrum (CLA) is a small, yet highly interconnected subcortical structure in the mammalian brain (Bruguier et al., 2020) whose precise function remains mysterious. Due to its widespread and extensive reciprocal connections with the cortex, the CLA has been proposed to underlie various high-level functions including salience detection (Atilgan et al., 2021; Remedios et al., 2014; Smith et al., 2019), attentional control (Goll et al., 2015; White et al., 2020), and consciousness (Crick and Koch, 2005; Smythies et al., 2014). Additionally, results from recent studies suggest that CLA may be involved in associative learning (Reus-garcía et al., 2020; Terem et al., 2020), coordination of slow-wave activity (Narikiyo et al., 2020; Norimoto et al., 2020), distractor suppression (Atlan et al., 2018; White et al., 2018, 2020), attentional set shifting (Fodoulian et al., 2020), task switching (Krimmel et al., 2019), and behavioral engagement (Atlan et al., 2021; Liu et al., 2019).

Although its function remains unclear, the anatomy of inputs to and outputs from the CLA in rodents has been well characterized in recent years (Atlan et al., 2016, 2018; Mathur, 2014; Wang et al., 2017; Zingg et al., 2018). CLA projects to most cortical areas with varying strengths. Projections are strongest to higher-order and associative cortical areas, and weakest to primary sensory and motor areas (Atlan et al., 2018; Wang et al., 2017; White et al., 2016; Zingg et al., 2018). Within the areas CLA projects most strongly, axons are found most densely in layer 2/3, followed by layers 6 and 5 (Wang et al., 2017). Whole-brain reconstructions of single CLA neurons have revealed that individual neurons project to an average of 21 cortical regions (Peng et al., 2021; Wang et al., 2022). In addition, CLA neurons are arranged topographically; projections from anterior CLA neurons are biased to innervate anterior cortical areas, while posterior CLA neurons project to posterior cortex (Wang et al., 2022). Slice electrophysiology experiments have demonstrated that CLA neurons can be categorized into several different types based on their intrinsic electrical properties (Chia et al., 2017; Graf et al., 2020); though it is unknown whether these correlate with differences in cortical projection targets.

Most cortical regions that CLA projects to form reciprocal connections, with a few notable exceptions that send few or no projections back to the CLA. For example, the anterior cingulate (ACA) both receives strong input from and projects back to CLA, whereas the retrosplenial cortex (RSP) receives substantial input from CLA but only sparsely projects back to CLA (Wang et al., 2017; Zingg et al., 2018). Work using brain slices found that CLA neurons projecting to ACA are less likely to receive input from sensorimotor regions, and vice-versa (Chia et al., 2020). Together these studies suggest that the connections between CLA and cortex may form distinct functional modules (see also Marriott et al., 2020).

Recently, several studies have used optogenetic stimulation of the CLA and recordings in frontal cortex *in vivo* to investigate the functional impact of claustro-cortical projections on the cortex (for a review, see Jackson et al., 2020). Two of these found a strong inhibitory effect in the prefrontal cortex (PFC) as a result of brief (5 ms) CLA stimulation (Jackson et al., 2018; Narikiyo et al., 2020). Since CLA neurons forming long-range projections to the cortex are glutamatergic, this CLA-stimulation-induced decrease in cortical neuron firing rates is likely due to feed-forward inhibition. Consistent with this, CLA axons were more likely to connect to cortical interneurons than to pyramidal neurons (Jackson et al., 2018). Another study also recorded in PFC but used different, prolonged CLA optogenetic stimulation patterns and found that a majority of frontal neurons increased in firing rate, rather than decreased (Fodoulian et al., 2020). In addition, a recent slice physiology study found that stimulation of CLA axons drove robust excitatory post-synaptic potentials in RSP pyramidal neurons (Brennan et al., 2021). Thus, the nature of CLA’s influence on cortex remains unresolved. Moreover, because the CLA projects to many cortical areas, it is unclear whether the CLA has the same impact on different regions, or whether CLA’s influence is more heterogeneous.

Here, we sought to broadly assess the functional impact of the CLA on multiple major cortical targets including PFC, secondary motor cortex (MOs), ACA, and RSP. To do this, we employed optogenetic stimulation of neurons expressed in the CLA in combination with multi-area Neuropixels recordings. We used two different Cre driver lines to selectively express channelrhodopsin in CLA cells. One line, Gnb4-Cre, has been very well characterized anatomically and expresses Cre mostly in the CLA (Peng et al., 2021; Wang et al., 2017). The other line, Esr2-Cre, expresses Cre more selectively in the CLA, but less completely, and in fewer cells overall than the Gnb4-Cre line (Wang et al., 2022).

We used Neuropixels probes (Jun et al., 2017) to simultaneously record spiking activity from 15,444 units across cortical layers 2/3, 5, and 6. In addition, by registering brains to the Allen Mouse Brain Common Coordinate Framework (CCF; Wang et al., 2020), we estimated the position of each recorded unit to within 100 μm. We find that the effect of CLA stimulation varies with different stimulation parameters; short pulses briefly activate putative inhibitory cells, while prolonged stimulation produces more excitation (see Supplementary Movies). The impact of CLA stimulation on cortex also varies across cortical areas and layers, and is not simply a blanket of inhibition. Indeed, many cortical neurons are robustly excited by CLA activity. These results suggest that the function of claustro-cortical projections is diverse, and demonstrates the need to study subpopulations of CLA neurons projecting to different cortical areas, as they may play distinct roles in brain function.

## RESULTS

### Optogenetic claustrum stimulation and Neuropixels recordings from distributed cortical areas

To characterize the impact of claustrum (CLA) on medial cortical areas, where its cells project densely (Figure 1A), we implanted Gnb4-IRES2-CreERT2;Ai32 mice with an optic fiber cannula above the left CLA (Figure 1B,C). Then, in awake, head-fixed mice that were free to move on a disc, we optogenetically stimulated CLA with different light patterns (Figure 1A, lower right) and used Neuropixels probes to record units from up to 3 simultaneous insertion locations across the anterior-posterior axis of the same, left hemisphere (Figure 1D,E). The four different stimulation patterns were as follows: a single 5 ms pulse, a series of 5 ms pulses at 20 Hz or 40 Hz for 500 ms (10 and 20 pulses, respectively), and a prolonged step of 500 ms duration. The 5 ms pulse is a similar stimulation pattern to previous studies (Jackson et al., 2018; Narikiyo et al., 2020), while the 500 ms stimulation reflects the observation that CLA cells can become active for prolonged periods of time (100-500 ms) in a behavioral context (Chevée et al., 2021; Ollerenshaw et al., 2021). On average, we estimate that we stimulated 30-75% of CLA volume (Figure S1).

**Figure 1:**
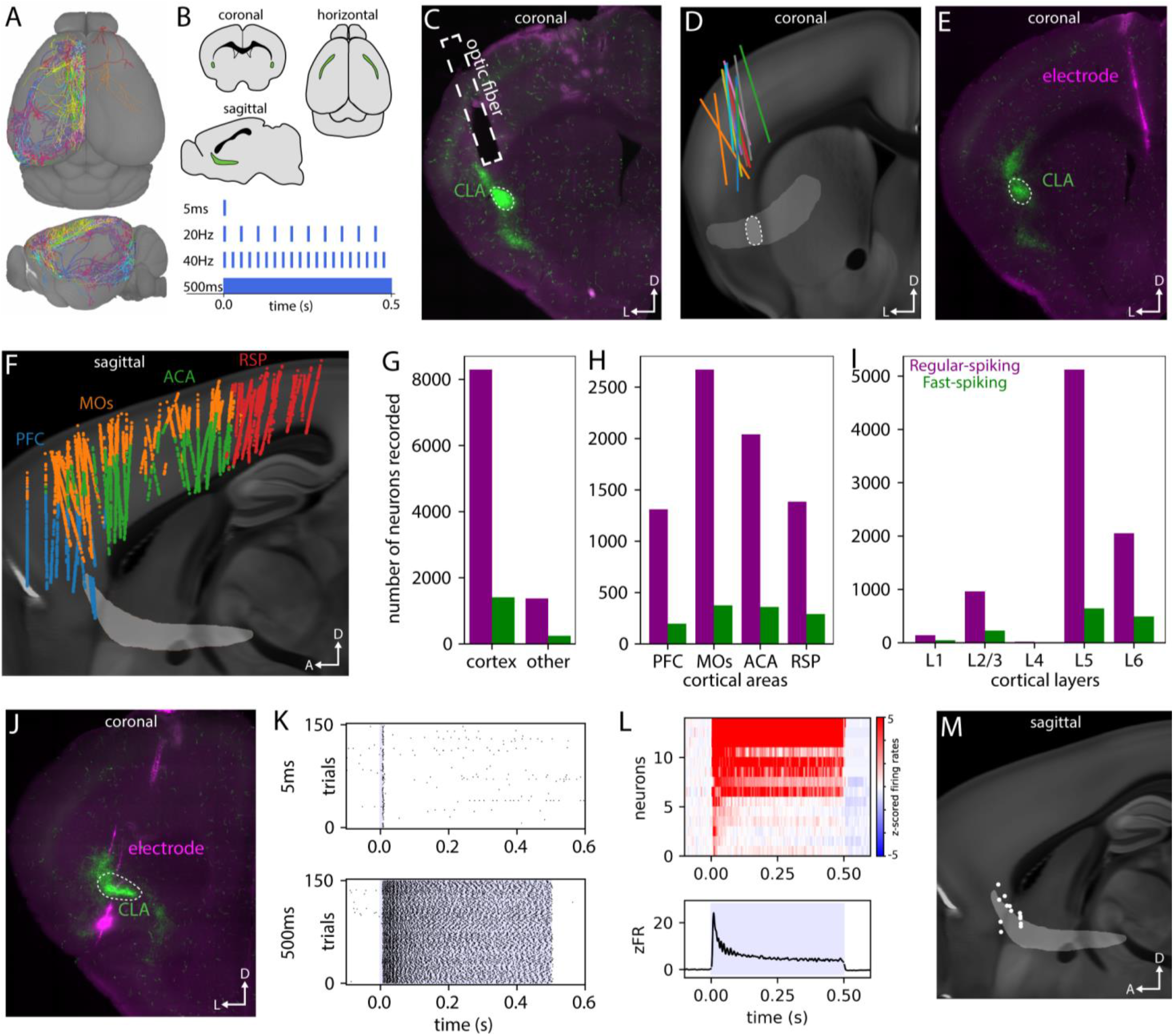
Optogenetic perturbation of claustrum and Neuropixels recordings in cortex. **A**, a subset of single claustrum (CLA) cell whole-brain projections, modified with permission from Peng et al., 2021. **B**, upper: coronal, horizontal, and sagittal views of the mouse brain with CLA highlighted in green. Lower: schematic of the four optogenetic stimulation patterns used to perturb the CLA. **C**, fluorescence microscopy image of a coronal section (dorsal=top, lateral=left) from a Gnb4-Ai32 mouse with implanted optic fiber above CLA. Green shows YFP fluorescence linked to channelrhodopsin, magenta shows autofluorescence in the red channel. **D**, reconstructed positions of implanted optic fibers in 13 Gnb4-Ai32 mice projected onto the Allen Mouse Brain Atlas. A 2-d projection of CLA across the entire extent of the brain is shown in light gray, with the location of CLA in this section highlighted in lighter gray. On average, we estimate that approximately 40% of CLA was illuminated across experiments (see Figure S1). For atlas alignment details see Methods. **E**, fluorescence microscopy image of a section from a Gnb4-Ai32 mouse showing a Neuropixels probe track in cortex. These images were aligned to the atlas to provide the location of recorded units. **F**, sagittal view (anterior=left, dorsal=top) of locations of recorded units from the primary cortical regions of interest: Prefrontal cortex (PFC; includes prelimbic (PL), infralimbic (ILA), and orbital (ORB) cortical areas), Secondary motor area (MOs), Anterior cingulate area (ACA), and Retrosplenial area (RSP). **G**, bar plot of the number of high-quality units recorded in cortex versus other brain regions. See figure S1 for further breakdown of non-cortical units. 85.9% of recorded neurons were in cortex, 85.2% of which were regular-spiking (RS, purple) and the remainder fast-spiking (FS, green) neurons. **H**, bar plot of the number of units recorded in each of the 4 main areas studied. See Figure S1 for further breakdown of units in other areas. **I**, bar plot of the number of units recorded across different cortical layers. **J**, fluorescence microscopy image of a successful CLA recording experiment. Neuropixels probe tracks (magenta) can be seen intersecting with CLA (green). **K**, raster plots from a putative CLA neuron responding to a single 5 ms optogenetic pulse (upper) or to a 500 ms sustained pulse (lower). **L**, upper, heatmap showing the z-scored responses of 14 putative opto-tagged CLA neurons to a 500 ms pulse. For details on CLA neuron opto-tagging criteria, see methods. Lower, average z-scored firing rates across the 14 CLA neurons. **M**, locations of the 14 opto-tagged CLA neurons on a sagittal view of the brain atlas.

In 92 experiments from 13 Gnb4-Ai32 mice, we recorded 9,658 units from cortical regions and 1,580 in subcortical areas (Figure 1F, S1). We focused our analyses on major target areas of CLA cell projections: Prefrontal cortex (PFC, combined units from Prelimbic, Infralimbic, and Orbital cortices), Secondary Motor cortex (MOs), Anterior Cingulate (ACA), and Retrosplenial (RSP) (Figure 1G). We recorded few neurons in layer 1 or layer 4 (208 total; most targeted regions were agranular); we therefore restricted analyses to layers 2/3, 5 and 6 (Figure 1H), leaving the following unit number totals per area: PFC: 1,428; MOs: 3,022; ACA: 2,347; RSP: 1,604. We divided units into two categories based on the width of their spike waveforms: regular-spiking (RS, putative excitatory, peak to trough ≥0.4 ms) and fast-spiking (FS, putative inhibitory, peak to trough <0.4 ms).

To validate optogenetic stimulation of CLA neurons, we targeted Neuropixels recordings to the CLA in ten experiments (Figure 1I-L). We recorded 14 units that were very likely to be opto-tagged CLA neurons based on their strong, low-latency activation and proximity to the CLA structure obtained from atlas registration (Figure 1K).

### Cortical cells can be excited or inhibited by claustrum stimulation in the Gnb4-Ai32 transgenic line

When optogenetically stimulating CLA, we observed a wider variety of cortical unit responses than in previous reports (example units, Figure 2A-C). Whereas previous studies observed primarily inhibitory responses in cortex, we found many units that were consistently excited, in addition to units that were inhibited across trials.

**Figure 2:**
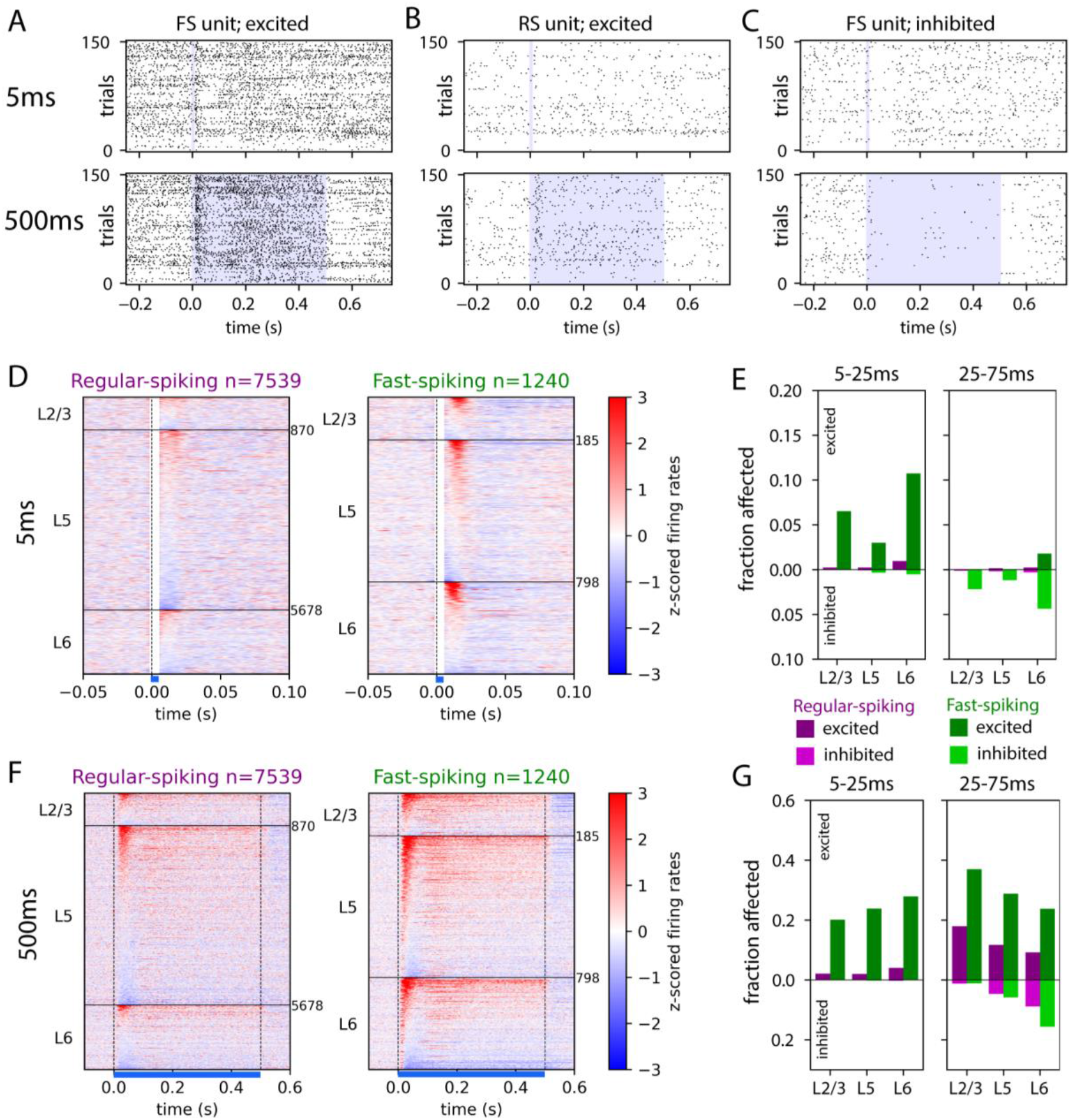
Cortical cells can be excited or inhibited by claustrum stimulation in the Gnb4-Ai32 transgenic line. **A-C**, raster plots from example units showing their responses to 5 ms (top row) and 500 ms (bottom). A and B show units that were excited by the 500 ms stimulation, and C shows a unit that was inhibited. Shaded area shows optogenetic stimulation duration. **D**, Heatmaps showing the average z-scored responses of regular-spiking (RS) units, left, and fast-spiking (FS) units, right, to a 5 ms optogenetic stimulation, across PFC, MOs, ACA, and RSP. Each row is one unit, sorted first by cortical layer and second by response magnitude. Cumulative number of units across layers shown on the right side of each panel. **E**, fraction of neurons significantly affected by the perturbation within two different time windows, separated by cortical layer. RS units are plotted in purple, FS units in green. The fraction of units with increased firing rates relative to baseline (excited) are plotted above 0, while units with decreased firing rates (inhibited) are plotted below 0. **F, G**, same as D, E, but for 500 ms sustained optogenetic stimulation. See Methods for details of how a unit was designated to be significantly affected.

In identical experiments with control mice that do not express channelrhodopsin, we observed no change in activity until an increase starting at approximately 100 ms (Figure S2), which could be due to heating of the tissue (Stujenske et al., 2015) or indirect photostimulation of the retina by the light source. Thus, to quantify the CLA-stimulation-specific responses, we restricted analysis of all four types of optogenetic stimulation parameters to an early (5-25 ms) and a late (25-75 ms) window following onset of optogenetic stimulation to exclude any light onset artifact and any later activity that could be due to other factors described above. To compute the fraction of units that significantly responded to CLA stimulation, we compared trial-by-trial spike counts in these early and late time windows versus baseline activity prior to stimulation (see Methods). In addition, for most analyses, we focused on trials when the animal was stationary (rather than locomoting); this corresponds to 83% of trials on average (Figure S2). Overall we observed elevated baseline firing rates in cortical neurons during locomotion compared to when mice remained stationary, and a smaller effect size of optogenetic stimulation, perhaps due to neurons already approaching their maximum firing rate (Figure S2).

Using a 5 ms pulse of light (as in previous studies), we observed significant excitation of primarily FS units in the 5-25 ms time window (6.1% of 1,176 FS units) and some inhibition of FS units in the 25-75 ms time window (2.4%), but very few RS units were either excited or inhibited (<0.5% of 7,225 RS units either excited or inhibited in either time window; Figure 2D,E). Next, using a 500 ms prolonged optogenetic stimulation, we found again that primarily FS units were excited in the early time window (24.6% of FS units versus 2.5% of RS units). However, in the later time window, we observed many more excited RS units (11.8%), suggesting that different patterns of CLA activation can produce different effects in its downstream cortical projection targets (Figure 2F,G; Supplemental Movies 1-2).

Our Neuropixels recordings spanned all cortical layers. Thus, we were able to assess the impact of CLA stimulation on different layers by dividing units into categories depending on their layer location. With both the 5 ms and 500 ms stimulation patterns, layer 6 contained the largest fraction of inhibited units for both FS (5 ms: 4.3%; 500 ms: 25.1%; 391 layer 6 FS units) and RS (5 ms: 0.3%; 500 ms: 8.8%; 1,653 layer 6 RS units), and layer 2/3 the fewest for both FS (5 ms: 2.2%; 500 ms: 1.1%; 184 layer 2/3 FS units) and RS (5 ms: 0.01%; 500 ms: 1.2%; 853 layer 2/3 RS units; Figure 2E,G). In contrast, layer 2/3 contained the largest fraction of excited units with 500 ms stimulation for both FS (37.0%) and RS (17.9%; Figure 2G).

Using 20 and 40 Hz pulse trains, the responses to each pulse resembled the responses to a single 5 ms pulse; primarily only FS units were activated (Figure S3). On average, units demonstrated adaptation to each successive pulse (Figure S3). This adaptation could be due to a combination of many factors, including properties of the opsin, adaptation of stimulated CLA neurons, or of the cortical neurons themselves. Interestingly, the opto-tagged CLA neurons that we identified displayed a slight facilitation with 20 Hz stimulation, and depression with the 40 Hz stimulation, suggesting that 20 Hz may be a more optimal frequency with which to stimulate the CLA.

### Neural responses to claustrum stimulation differ based on cortical area

Next, we investigated the dependence of responses on the cortical target area by dividing units into categories based on area, regardless of layer. In response to the 5ms pulse, a small and similar proportion of FS units were activated in PFC, MOs, and ACA in the early, 5-25 ms, time window (PFC: 5.2%; MOs: 10.0%; ACA: 5.7%), but fewer neurons were activated in RSP (1.9% Figure 3B). In the late, 25-75 ms, window, most inhibited units were observed in area MOs (6.7%).

**Figure 3:**
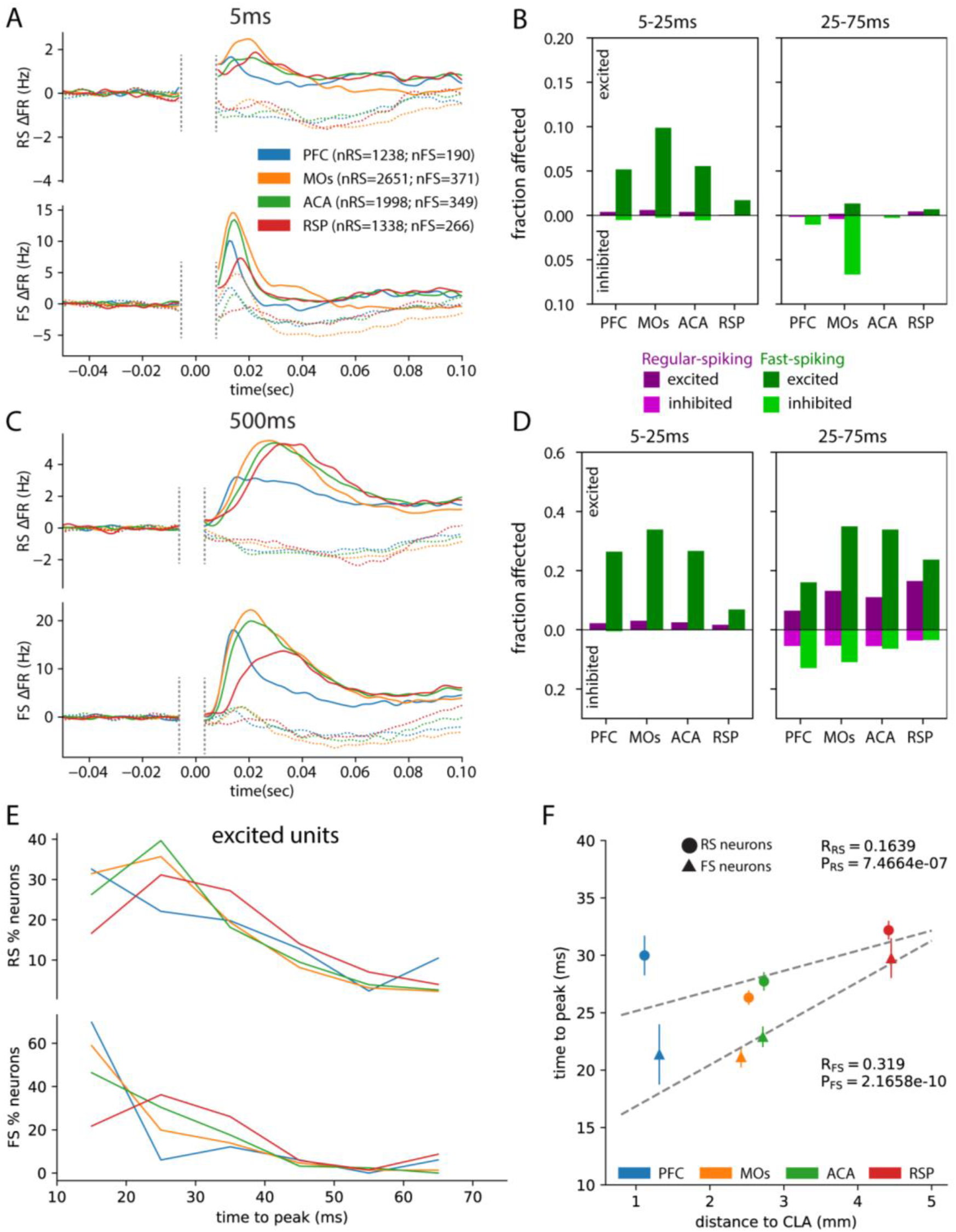
Neural responses to claustrum stimulation in Gnb4-Ai32 mice differ based on cortical area. **A**, Average baseline-subtracted PSTHs of all recorded units in different cortical areas (PFC, blue; MOs, orange; ACA, green; RSP, red) in response to a 5 ms optogenetic stimulation. Upper, RS; lower, FS; excited units, solid lines; inhibited units, dotted lines. The area around t=0 denoted by vertical dashed lines is blanked out because of the presence of an optogenetic onset artifact in many of the experiments (see Methods). **B**, fraction of neurons significantly affected by the 5 ms stimulation, separated by cortical area. **C**, **D**, same as A, B, but for the 500 ms sustained optogenetic stimulation. See Methods for details of how a unit was designated to be significantly affected. **E**, histograms of time to peak firing rate for RS (upper) and FS (lower) neurons that were significantly excited in response to the 500 ms sustained stimulation, across the different cortical areas. **F**, Average time to peak for each area and cell type versus distance to CLA; each point shows the average and variation across units from a given area and cell type combination. The time to peak of both RS and FS units significantly correlated with their distance from CLA (Pearson correlation run across all FS or RS units). Error bars SEM.

In response to the 500 ms stimulation, the pattern of activation of cortical units was similar in the early time window, but stronger (Figure 3D). Relatively few RS units were excited (PFC: 2.0%; MOs: 3.0%; ACA: 2.5%; RSP: 1.7%), while many FS units were excited (PFC: 26.3%; MOs: 34.0%; ACA: 26.9%; RSP: 7.1%). In the late time window, different areas showed consistent differences in the relative fraction of excitatory and inhibitory responses. PFC and MOs contained the largest fraction of inhibited RS and FS units (PFC RS: 5.6%, FS: 13.2%; MOs RS: 5.4%, FS: 11.1%), while RSP contained the largest fraction of excited RS units (16.3%) and the smallest fraction of inhibited units (RS: 3.7%, FS: 3.4%; Figure 3D). This demonstrates that the net effect of prolonged CLA stimulation on the cortex depends on cortical area. In PFC, the proportion of excited and inhibited units was approximately equal (RS: 6.1% excited, 5.8% inhibited; FS: 14.7% excited, 13.2% inhibited), while in RSP, the majority of affected units were excited (RS: 16.3% excited, 3.7% inhibited; FS: 23.7% excited, 3.4% inhibited).

Interestingly, we noticed that the time to peak of population responses from each of these cortical areas was different, particularly for FS units, with frontal areas peaking earlier on average (PFC: 21.36 ms; MOs: 21.15 ms) than more posterior areas (ACA: 22.90 ms; RSP: 29.75 ms; Figure 3A,C). To examine this in more detail, we computed the distribution of time to peak (within the first 75 ms) across each of the cortical areas for the subset of units that were excited by CLA stimulation. This demonstrated that units in frontal areas tend to reach their peak firing rate more quickly than the others following CLA stimulation (Figure 3E; Supplemental Movies 1-2). We then used the CCF coordinates of each unit to find its Euclidean distance from the nearest point of CLA, plotted time to peak versus distance to CLA for each unit, and computed the Pearson correlation across RS and FS units separately (Figure 3F). We found a significant correlation between excited units’ distance from CLA and time to peak firing rate, consistent with the path of CLA axons that travel first to frontal regions then travel backward toward posterior areas like RSP, with a slope of 3.61 ms/mm for FS units and 1.75 ms/mm for RS units. If the time to peak activation were dependent on the CLA axon conduction velocity alone, we would expect these slopes to be identical. The difference may reflect that FS neurons tend to be recruited most quickly in frontal regions and delay activation of RS neurons, differences in synaptic strength between CLA axons and FS and RS neurons across areas, and differences in intrinsic properties of the neurons themselves. We also computed these measures for units inhibited by CLA stimulation, and found a similar positive correlation, though it was only significant for FS units (Figure S3).

### Replication of claustrum stimulation experiment in the Esr2-IRES-Cre;Ai32 line

Though the Gnb4-IRES2-CreERT2 transgenic line expresses Cre largely specifically to CLA, there are still some non-CLA cells that express Cre, particularly in the deep layers of the overlying insular cortex (Peng et al., 2021; Wang et al., 2017). Therefore, we sought to replicate the above experiments in a different transgenic mouse line, Esr2-IRES2-Cre, which more selectively expresses Cre in CLA than the Gnb4 line, albeit less completely (Figure 4A; Wang et al., 2022). We crossed the Esr2-IRES-Cre line to the channelrhodopsin reporter line, Ai32, and implanted optic fiber cannula as with the Gnb4-Ai32 mice (Figure 4A,B). In these mice, we validated optogenetic stimulation with CLA-targeted recordings (Figure 4C,D) as we did with the Gnb4-Ai32 mice. In cortex, we recorded 7,043 units across PFC, MOs, ACA, and RSP (Figure 4E-H). Overall, the responses of cortical units to CLA perturbation in the Esr2-Ai32 line were weaker than those in the Gnb4 line (Figure 4I-L). 5 ms stimulation evoked little activity (see Figure S4); therefore, we here focus on the responses to 500 ms stimulation. FS cells were more likely to be recruited in both the early (RS: 0.34%, FS: 5.24%) and late (RS:4.04%, FS: 9.64%) analysis windows. Focusing on the late, 25-75 ms time window, Layer 6 contained the highest fraction of inhibited units (RS: 5.60%, FS: 14.42%), while layer 2/3 contained the largest fraction of excited units (RS: 8.94%, FS: 14.20%; Figure 4K). Excited RS units were most common in RSP (7.17%; Figure 4L), and least common in frontal areas (PFC: 2.02%; MOs: 2.89%). Interestingly, the fraction of inhibited units also tended to increase from anterior (PFC RS: 1.82%, FS: 7.69%) to posterior (RSP RS: 4.74%, FS: 9.91%), which is the opposite of Gnb4-Ai32 experiments. This could suggest that Esr2+ CLA cells tend to project more strongly to RSP relative to PFC. Nonethess, overall, we see a largely similar pattern of activity in the cortex when using two different transgenic mouse lines to express channelrhodopsin in CLA (also see Supplemental Movies 3 and 4). The smaller overall number of neurons stimulated with the Esr2 line is consistent with the weaker overall responses evoked with this line compared to Gnb4, which labels more CLA neurons.

**Figure 4:**
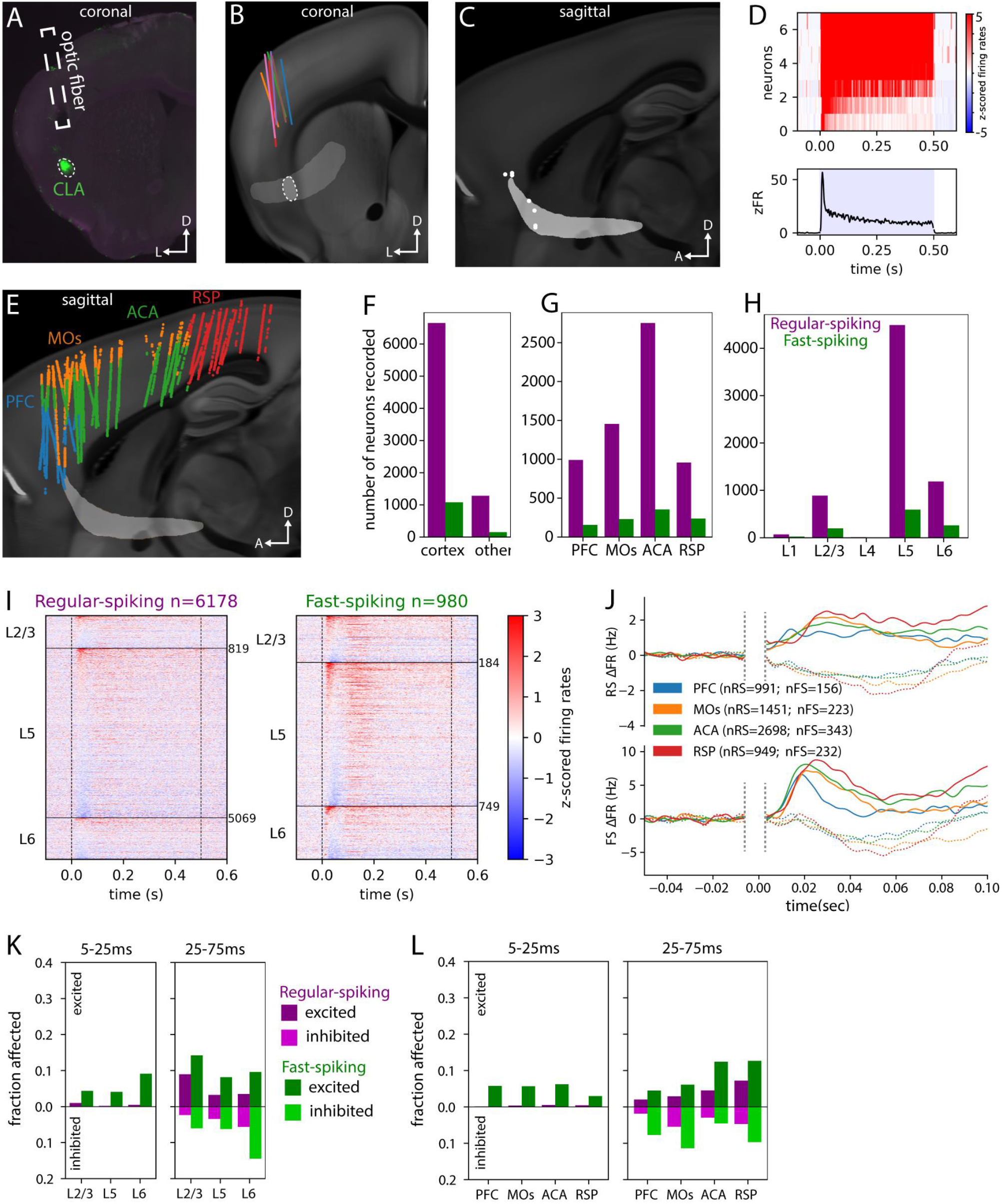
Cortical responses to claustrum stimulation in Esr2-Ai32 transgenic line. **A,** histology image (dorsal=top, lateral=left) showing the location of an optic fiber implant above CLA in an Esr2-IRES2-Cre;Ai32 mouse. **B**, reconstructed positions of implanted optic fibers in 7 Esr2-Ai32 mice projected onto a coronal section of the CCF. **C**, locations of the 7 opto-tagged CLA neurons recorded from Esr2-Ai32 mice on a sagittal view of the CCF (anterior=left, dorsal=top). **D**, upper, heatmap showing the z-scored responses of the 7 putatzve opto-tagged CLA neurons. Lower, average z-scored firing rates across the 7 CLA neurons. **E**, sagittal view of locations of recorded units from the primary cortical regions of interest: Prefrontal cortex (PFC; includes prelimbic (PL), infralimbic (ILA), and orbital (ORB) cortical areas), Secondary motor area (MOs), Anterior cingulate area (ACA), and Retrosplenial area (RSP). **F**, number of units recorded from cortex versus other brain structures. **G**, number of units recorded across the 4 main brain regions studied: PFC, MOs, ACA, and RSP. **H**, number of units recorded across different cortical layers. **I**, heatmap showing the average z-scored responses of regular-spiking (RS) units, left, and fast-spiking (FS) units, right, to a 500 ms optogenetic stimulation, across PFC, MOs, ACA, and RSP. Each row is one unit, sorted first by cortical layer and second by response magnitude. **J**, average baseline-subtracted PSTHs of units in different cortical areas in response to a 500 ms optogenetic stimulation. Upper, RS; lower, FS; excited units, solid lines; inhibited units, dotted lines. Vertical dotted lines denote data not included in analysis due to an optogenetic artifact present in some recordings. **K**, fraction of neurons significantly affected by the perturbation within different time windows, separated by cortical layer. RS units are plotted in purple, FS units in green. The fraction of units with increased firing rates relative to baseline are plotted above 0, while units with decreased firing rates are plotted below 0. **L**, fraction of neurons significantly affected by the 500 ms stimulation by cortical area. See Methods for details of how a unit was designated to be significantly affected.

### Functional clustering reveals cortical cells with distinct responses to claustrum stimulation

To characterize the diversity of cortical responses to CLA stimulation, we clustered 15,444 neurons from both Gnb4 and Esr2 lines based on their trial-averaged response to the 500 ms stimulation. Clustering was restricted to 5-75 ms post-stimulation to avoid possible contamination due to thermal or visually evoked activity (Figure S2). Pairwise cosine similarity of z-scored responses between all pairs of responsive neurons (n=3,199) was computed and used as features to cluster on (Figure 5A). The feature vectors were fit to a Gaussian Mixture Model to obtain 23 clusters, which were further merged to obtain a final 9 different response types (see Methods, Figure S5A). Individual neural responses to the 500 ms as well as to the 5 ms stimulation data within all 9 clusters were found to have a high correlation with the mean response of the parent clusters (but not to the other 8 clusters), indicating good quality of clustering (Figure 5B, S5B, C).

**Figure 5:**
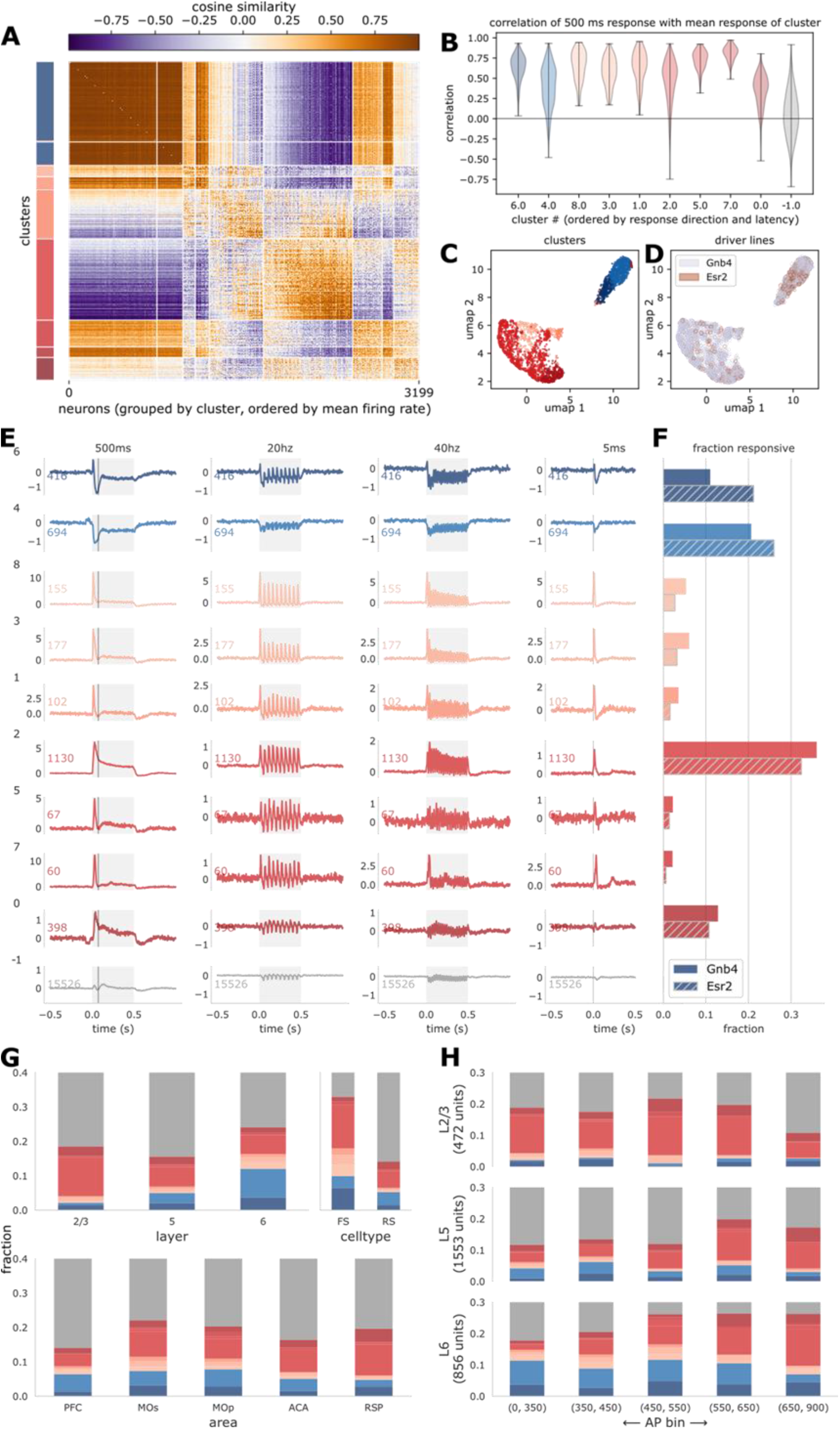
Functional clustering of neurons based on mean response to 500 ms CLA stimulation. **A.** Cosine similarity matrix of mean initial response (0 − 75 ms) of all 3,199 responsive neurons to the 500 ms stimulation from both Gnb4 and the Esr2 transgenic lines. The similarity vectors (rows of the matrix) were clustered using a Gaussian Mixture Model. Neurons are grouped into 9 clusters (color bar on the left) and ordered by mean firing rate within clusters for visualization. Clusters that were predominantly inhibited are colored in shades of blue, and those that were excited are in shades of red. Darker shades correspond to later peak (or minimum) responses. Colors are consistent across this and related figures. **B.** Distribution of correlation of individual neural responses with mean response of their cluster. High correlation values and distributions largely above 0 indicate a high quality of resultant clusters. Similar metric computed for 15,526 unresponsive neurons shown in grey for reference (on average zero correlation with mean response of unresponsive neurons). **C.** A 2D projection of the similarity vector for each neuron obtained using UMAP, colored by cluster identity. Two major classes of response types were observed, corresponding to predominant inhibition (blue) and excitation (red), with further clusters within each class. **D.** Same 2D projection as in (C) colored by the optogenetic driver line. Esr2 and Gnb4 are well-mixed, indicating that they drive similar populations in similar ways. **E.** Mean responses of each cluster to all stimuli. Numbers to the left of the plots indicate cluster number. Annotations with the same color as the cluster indicate number of neurons in the cluster. Blue are inhibited and red are excited clusters. Shade indicates latency to the peak (or minimum) response. **F.** Fraction of responsive neurons that belong to each cluster. Overall Esr2 (hatched) and Gnb4 (solid) drivers result in a similar distribution of response types. Esr2 has a slight preference for inhibition of cortical neurons compared to Gnb4. **G.** Distribution of different response types across layers, areas or cell types. **H.** Distribution of response types along the anterior-posterior axis. Units indicated in the AP bins correspond to 10μm voxels in CCF space. Bins were chosen to contain approximately equal numbers of neurons.

A 2D UMAP projection of the feature vectors revealed two well-separated classes of neurons, corresponding to inhibited (~1,000 neurons) and excited (~2,000 neurons) clusters (Figure 5C). Both 2D projections also showed that the responses driven by Esr2 and Gnb4 drivers were well-mixed (Figure 5D, Figures S5D), indicating that the two driver lines evoke similar response patterns in the cortex including both excitatory and inhibitory effects.

We identified several different response profiles (Figure 5E), including neuron classes that are only excited or inhibited, or those that show a transient excitation followed by inhibition and vice versa. The post-stimulation response latency varied across these clusters from 10 ms to 55 ms. The relative fractions of neurons belonging to each cluster were similar across Gnb4 and Esr2 (Figure 5F), further underscoring the similarity between the two lines. For instance, in both lines, about one third of all cortical cells were of the ‘sustained excitation type’ (excitatory cluster 2; Figure 5F). We did observe, however, that there was a slight bias for inhibitory responses in Esr2 line.

Although the neurons were clustered based on responses to the 500 ms stimulus, they showed similar trends in responses to other stimulation patterns (Figure 5E, S6). For instance, the clusters that were net inhibited by the 500 ms stimulus were also inhibited, on average, for the 20 and 40 Hz pulse trains. Interestingly, these inhibited clusters also showed evidence of a short-latency excitatory response that was quickly followed by a suppression of spiking. Most clusters showed distinct responses to each pulse of the 20 Hz pulse train, but the individual responses are less clear at 40 Hz. Thus, cortical neurons are unable to follow modulations of firing rate in the CLA occurring beyond a threshold frequency between 20 and 40 Hz.

RS versus FS cells showed some bias across clusters (Figure 5G). In particular, early responding excited clusters (“salmon-colored” clusters; clusters #8, 3, 1) showed a rapid decrease in firing rate (below the baseline in one case) following the initial peak. These neurons were enriched among the FS population (Figure 5G), likely reflecting interneurons that are initially excited by direct input from the CLA followed by an inhibition. The initial peak followed by inhibition below baseline is more evident in the 5 ms responses of these three clusters (Figure S6).

Across layers, we found layer 6 to have the highest fraction of inhibited neurons, and layer 2/3 the most delayed responders. We did not observe striking differences across areas, except RSP, which was enriched for late responders. In general, the fraction of late responders increased going from anterior to posterior neurons, especially in deep layers (Figure 5H), consistent with the anatomical distance along axons projecting from CLA.

### Claustrum stimulation does not synchronize cortical areas in the gamma frequency band

One hypothesis of CLA function is that, with its widespread cortical projections, it can coordinate or synchronize cortical areas and act as a conductor of those areas (Crick & Koch 2005). To test this hypothesis, we used signals from the local field potential band (0.1-500 Hz sampled at 2.5 kHz), recorded from the same Neuropixels probes as the spiking activity, and computed metrics of local field potential phase synchronization, spike-field coupling, and spike-spike coupling. Due to the onset and offset transients induced by the optogenetic stimulation, the local field potential responses in the 20 Hz and 40 Hz were likely contaminated by these artifacts, so we focused on the responses to 500 ms stimulation.

First, we computed the oscillatory power in the local field potential (LFP) signals in the baseline compared to the 500 ms stimulation (25-475 ms time window) and averaged these power spectra across channels within each cortical area for each experiment (Figure 6A). We found a significant increase in power in the high gamma band, defined here as 65-80 Hz, in all four examined cortical areas (Figure 6B; paired t-tests with Benjamini-Hochberg false discovery rate correction). We also observed this increase in power in Esr2 mice, but not in control wildtype mice in which the identical optical fiber delivered the same light stimulus (Figure S7).

**Figure 6:**
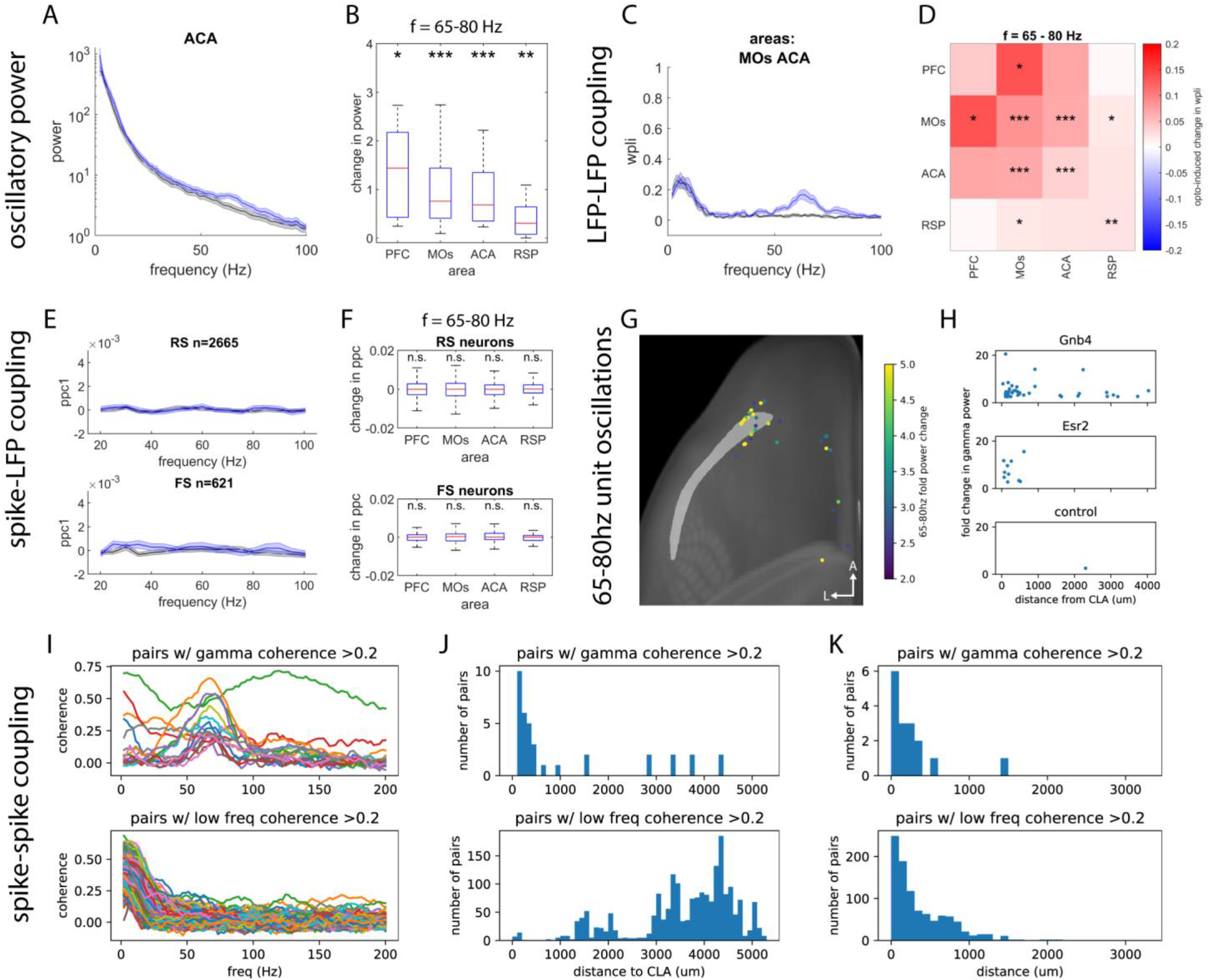
Measures of functional connectivity. **A,** example power spectrum obtained from local field potentials recorded in ACA, averaged across 20 experiments. The black line shows the power spectrum prior to, and the blue line during optogenetic stimulation. **B,** summary change in power in the 65-80 Hz gamma band in different cortical areas across all experiments. Individual datapoints are averages of the power during minus the power prior to CLA stimulation from each experiment. Power in the gamma band significantly increased in all 4 areas. **C,** example phase synchronization spectrum obtained from pairs of LFP signals recorded in MOs and ACA. Black shows the debiased phase lag index prior to optogenetic stimuli, and blue shows the debiased weighted phase-lag index (WPLI, see text and Methods for details) during optogenetic stimuli. **D,** LFP-LFP connectivity heatmap, showing the strength and direction of CLA stimulation-induced change in coupling between pairs of areas. Red indicates an increase, and blue a decrease. Matrix is symmetric. **E**, Average pairwise phase consistency spectra across included recordings. **F**, change in PPC across different areas and cell types. We did not observe a significant CLA-stimulation-induced change in spike-field coupling in the gamma band. **G**, horizontal view of the brain atlas (anterior top, lateral left) with CLA shown in light grey and the position of single units with a large CLA-stimulation-induced change in 65-80 Hz power. The fold change of power indicated with color: yellow for a large change, blue for a smaller change. **H,** Plot of the fold change in units’ 65-80 Hz power versus their distance from CLA, across different transgenic lines. In both Gnb4 and Esr2 lines, most units with a large increase in gamma power are located closer to CLA. **I**, Coherence spectra of pairs of units. Upper, pairs with a peak in the gamma range; lower, pairs with a peak in the lower frequency range (<10 Hz). **J**, Histograms comparing the distribution of the unit pairs’ positions relative to CLA. Most pairs with a gamma peak are located close to CLA, while pairs with low frequency coherence are more widespread. **K**, Histograms comparing the distance between these units. Most unit pairs with gamma coherence are located close to each other. This is also true for units with low frequency coherence, but there are many more that are >500um away from each other. Significance indicators: *p<0.05, **p<0.01, ***p<0.001, paired t-tests with Benjamini-Hochberg false discovery rate correction. Shaded error bars SEM; box plots show medians, quartiles, and 95% confidence intervals.

We also investigated the oscillatory characteristics of the units themselves. We computed the power spectra of the units’ spike trains across trials. A handful of units had increased gamma oscillations during CLA stimulation, and these units tended to be located near the CLA; consequently these are likely CLA neurons (Figure 6G,H). Thus, during the 500 ms stimulation, CLA neuron spiking has a tendency to oscillate in the gamma band, consistent with increase in LFP power measured in the cortex in this frequency band.

Next, we examined phase synchronization between the signals recorded on pairs of channels in the same or in different areas. To quantify this, we used the debiased weighted phase lag index (WPLI), which reduces the impact of volume conduction compared to the traditional coherence measure, and is bounded between 0 and 1; 0 indicates signals with no phase relationship and 1 indicates signals perfectly in phase (Oostenveld et al., 2011; Vinck et al., 2011). Averaging across pairs of channels within each combination of areas for each experiment, we found significant increases in phase synchronization caused by the 500 ms stimulation in the same gamma band where we observed increases in power (65-80 Hz; Figure 6C,D). We did not observe such an increase in phase synchronization between any area-pairs in either the Esr2 or control mice (Figure S7). In the Esr2 mice, this could be due to the much weaker sustained responses to optogenetic stimulation compared to Gnb4 mice.

LFP signals are thought to primarily reflect presynaptic activity. Thus, the increases in power and synchronization we observed could be largely due to the synchronized outputs from CLA. To assess whether the spiking activity within an area synchronized to these oscillations, we computed the pairwise phase consistency, a metric of spike-field coupling; a value above zero indicates spikes fired at a consistent phase in a particular frequency band (Vinck et al., 2010, 2012). We did not find a significant change in pairwise phase consistency in either any of the examined areas or RS vs. FS unit types (Figure 6E,F). This indicates that under these conditions, gamma oscillations in the CLA do not entrain downstream neurons in cortex.

If the spikes and LFP were not synchronizing in cortical areas in the gamma band, perhaps the increase in gamma power we observed was a result of the direct stimulation causing the local CLA circuit to oscillate. We computed the oscillatory power in the gamma band (65-80 Hz) of the spike trains from all recorded neurons both prior to and during the 500 ms optogenetic stimulation and selected those units with an absolute change in gamma power greater than 2.5-fold. These units were almost all located in or near CLA (Figure 6G,H), and the units we identified as opto-tagged CLA neurons in both Gnb4 and Esr2 lines tended to have a large increase in gamma power (mean fold change in Gnb4: 4.24±1.39; Esr2: 6.19±1.70; SEM).

We also examined unit-unit synchronization to assess between-area coordination. To do this we computed the coherence between pairs of cortical neurons. This measure was biased for units with higher firing rates (Figure S7). We observed unit pairs with low frequency (<10 Hz) coherence and pairs with gamma coherence (Figure 6I). Whereas unit pairs with low frequency coherence were broadly distributed throughout the brain, pairs with gamma coherence were located in the CLA (Figure 6J).

Together, these results suggest that the strong optogenetic activation of a large portion of the CLA causes the local circuit in the CLA to oscillate in the gamma range; however, this oscillation does not appear to entrain neurons in downstream cortical areas.

## DISCUSSION

Our results, based on recording from 15,444 cortical units in 13 Gnb4-Ai32 and 7 Esr2-Ai32 mice, demonstrate that the effect of claustrum (CLA) input on cortex is not always suppressive but can also be excitatory. Moreover, the effects vary depending on cortical area, layer, cell type, and CLA activation pattern (see the four supplementary movies).

In common with other reports (Jackson et al., 2018; Narikiyo et al., 2020), we find that a 5 ms short pulse of light activation of Gnb4+ CLA neurons with ChR2 triggers widespread activity in cortical interneurons and, to a much smaller extent, in cortical excitatory neurons. Across all recorded cortical areas, fast-spiking (FS; putative inhibitory) cells are 15 times more likely than regular-spiking (RS; putative excitatory) to be recruited within 25 ms of brief CLA stimulation (RS: 0.40%, FS: 6.12%, left of Figure 2E), consistent with Jackson et al. (2018) and Narikiyo et al. (2020).

In a later time window (25-75 ms), there is relatively little activation of either FS or RS units (RS: 0.15%; FS: 0.60%, right of Figure 2E). Despite this strong recruitment of putative inhibitory interneurons with the 5 ms activation of CLA we observe little reduction in firing activity of any type of cortical cells, particularly in RS cells (early time window: RS: 0.04%, FS: 0.34%; late time window: RS: 0.19%, FS: 2.38%). This lack of profound inhibition of excitatory cortical cells is different from what was reported by Jackson et al (2018) and Narikiyo et al (2020); this could be explained by a number of methodological differences that we discuss further below.

Prolonged 500 ms stimulation of Gnb4+ CLA neurons causes a large proportion of neurons to be activated in the cortex (see Supplemental Movies 1-2). In the first 25 ms following the onset of stimulation, FS cells are 10 times more likely than RS cells to be significantly activated (RS: 2.46%, FS: 24.6%, left of Figure 2G). Interestingly, in the later, 25-75 ms window, many more cortical RS cells are activated, such that FS neurons are only 2.4 times more likely to be activated (RS: 11.8%, FS: 28.4%, right of Figure 2G). This stimulation also suppresses many cortical neurons in the late window (RS: 5.18%, FS: 8.3%), though not as many as it excites. The observation that many RS and FS cortical cells are excited, while some others are inhibited, is consistent with a previous study that also used prolonged optogenetic stimulation of CLA (Fodoulian et al., 2020).

### Area- and layer-specificity of effect of CLA on cortical neurons

A novel observation from our experiments is that the balance between the proportion of excited versus inhibited neurons differed depending on cortical area and layer, particularly 25-75 ms following the onset of the prolonged CLA stimulation protocol (see Figure 3D). In frontal cortical areas, we observed the most inhibited neurons (PFC: RS: 5.6%, FS: 13.2%) and a similar proportion of excited neurons (PFC: RS: 6.1%, FS: 14.7%), while in posterior cortical areas, relatively few were inhibited (RSP: RS: 3.7%, FS: 3.4%) and more were excited (RSP: RS: 16.3%, 23.7%). These differences suggest that the function of projections from CLA to various cortical areas may be distinct.

The effect of CLA stimulation on RSP appears to be quite different from the other areas, with fewer neurons inhibited. RSP is also unique among the other cortical areas studied here because unlike PFC, ACA, and MOs, it does not send strong projections back to the CLA (Wang et al., 2017; Zingg et al., 2018). RSP is implicated in a variety of functions including spatial navigation, learning, and memory, and the default mode network (Alexander and Nitz, 2015, 2017; Keshavarzi et al., 2021; Miller et al., 2019; De Sousa et al., 2019; Vann et al., 2009; Whitesell et al., 2021). In addition, dissociative drugs cause an increase in 1-3 Hz oscillatory power in L5 of RSP (Vesuna et al., 2020), suggesting RSP is involved in regulating or maintaining conscious states. Given the excitatory influence of CLA on RSP and the lack of RSP projections back to CLA, the CLA-RSP projection may act as more of a driver or output pathway than the CLA projections to other cortical areas. This suggests that the CLA-RSP pathway should be studied in more detail moving forward, as it appears to be unique compared to the CLA-PFC or CLA-ACA projections that have been most studied to date.

We also found that the effect of CLA stimulation on the cortex depended on layer. Focusing on the cortical response 25-75 ms after the onset of 500 ms stimulation (see Figure 2G), a similar proportion of neurons were excited (L6: RS: 9.2%, FS: 23.8%) and inhibited (L6: RS: 8.8%, FS: 15.6%) in deeper layers, while most affected neurons in superficial layers were excited (L2/3: RS: 17.9%, FS: 37.0%) versus inhibited (L2/3: RS: 1.2%, FS: 1.1%). Since layer 2/3 neurons tend to project within the cortex, while neurons in deeper layers often project to thalamus or other subcortical areas, this could indicate that CLA input promotes intracortical processing, and reduces cortical-subcortical interactions.

### Replication of results in Esr2-Ai32 transgenic line

When using the Esr2-Ai32 transgenic line to express ChR2 more selectively, but in fewer CLA neurons, we find that the major results from the Gnb4-Ai32 experiments are recapitulated. While there are few responses overall to 5 ms stimulation (<1% of all neurons excited or inhibited, amounting to 14/7043 units, see Figure S4), we observe that FS units are 15 times more likely to be significantly activated (5.24%) than RS neurons (0.34%) within the first 25 ms of the prolonged 500 ms CLA stimulation (left of Figure 4K), compared to 10 times more likely in Gnb4-Ai32 experiments. In the later time window of 25-75 ms, more units of both types are activated (RS: 4.0%, FS: 9.6%, right of Figure 4K), and the ratio of FS to RS activation is 2.39, very close to that of Gnb4-Ai32 experiments in the same stimulation and analysis window, which was 2.40.

In addition, the area- and layer-dependence of the effect of CLA input is similar between Esr2-Ai32 and Gnb4-Ai32 experiments, especially in the 25-75 ms window during 500 ms stimulation (see Figure 4K and Supplemental Movies 3-4). Superficial layers tend to have more excited (L2/3 RS: 8.9%, FS: 14.2%) than inhibited (L2/3 RS: 2.3%, FS: 6.0%) neurons, while deeper layers tend to have more inhibited (L6 RS: 5.6%%, FS: 14.4%) than excited (L6 RS: 3.5%, FS: 9.6%) neurons. Interestingly, while we see a similar trend for increasing excitation in posterior (RSP RS: 7.2%, FS: 12.5%) versus anterior (PFC RS: 2.0%, FS: 4.5%) cortical areas, we do not observe the accompanying decrease in inhibited cells (see Figure 4L). In fact, we see the opposite – the proportion of inhibited cells also tends to increase from anterior (PFC RS: 1.8%, FS: 7.7%) to posterior (RSP RS: 4.7%, FS: 9.9%). This could indicate that Esr2+ CLA neurons are more likely to innervate RSP than PFC. This would be consistent with the finding that neurons in what some refer to as the CLA “core”, which is likely to correspond with the location of these Esr2+ neurons, project strongly to RSP (Marriott et al., 2020; Wang et al. 2022).

Finally, when clustering together the cortical unit responses from both lines, we did not observe a strong bias of any functional cluster for belonging to a particular transgenic line (Figure 5). The main difference was a slightly higher prevalence of excitatory responses in Gnb4+ stimulation experiments. This suggests that though the overall proportions of cells affected somewhat differ depending on whether Gnb4+ or Esr2+ cells are stimulated, the firing patterns they produce in the cortex are largely similar.

### Differences with previous studies

The most likely explanations for discrepancies between our results and those of previous studies include different CLA cell populations being stimulated, recording methods, and/or effective strength of the stimulation. In terms of the cell populations, the three previous studies that include CLA stimulation and neural recordings in the cortex used three different methods for expressing channelrhodpsin in CLA (Fodoulian et al., 2020, Vglut2-Cre mouse with AAV-DIO-ChR2-YFP injection in CLA; Jackson et al., 2018, AAVretro-syn-Cre in PFC, AAV5-DIO-ChR2-YFP in CLA; Narikiyo et al., 2020, Cla-Cre mouse with AAV-EF1a-DIO-ChR2-EYFP injection in CLA). One reason for these differences is that the mouse CLA is difficult to isolate, and its exact cellular and anatomical definition remains unsettled, so there is not one agreed upon operational method for labeling CLA neurons (Erwin et al., 2021; Mathur et al., 2009; Wang et al., 2022). It is quite likely that these different molecular targeting methods resulted in different populations of CLA or non-CLA neurons expressing ChR2, which when stimulated could produce different effects in the cortex. For example, while Jackson et al. (2018) found that claustrum neurons innervated layer 6 of PFC more strongly than layer 2/3, other studies have found the opposite pattern (Fodoulian et al., 2020; Wang et al., 2017; Wang et al. 2022), which may explain differences between their findings of a primarily inhibitory effect, and ours and those of Fodoulian et al. (2020) of a primarily excitatory effect.

On the other hand, because of these differences, the commonalities between previous results and ours may be even more convincing. Our results using two different transgenic lines also give us some insight – as discussed above, we observe a smaller effect of Esr2+ CLA neuron activation on the cortex compared to Gnb4+, though most other findings relating to area, layer, and cell type dependence are similar. It is important to note that not all Gnb4+ cells are located in the CLA; some are in layer 6 of the adjacent insular and gustatory cortices (Wang et al. 2022). Thus, it is possible that in our Gnb4-Ai32 line experiments, some portion of the effect of CLA stimulation may also be due to stimulation of these nearby cortical layer 6 cells. Esr2+ cells are more spatially restricted to CLA, but less completely label the structure, and there are fewer of them overall than Gnb4+ cells (Wang et al. 2022). Together, though Esr2+ and Gnb4+ cells likely reflect somewhat different populations, the similarity in results strongly suggests that the trends we see across area, layer, and cell types in cortex are attributable to CLA neuron input.

The previous studies also used different recording methods – while all employed electrophysiology, the probes used were different. Jackson et al (2018) used tungsten electrodes, Narikiyo et al (2020) and Fodoulian et al (2020) used 32-channel silicon probes, and we used 384-channel Neuropixels probes. Due to their higher density of electrodes, Neuropixels probes can detect many more units than a single tungsten electrode or even a 32-channel probe (Jun et al., 2017). Thus, our study is based on 15,444 units compared to on the order of 100 units for the previous three studies, and is also less likely to be biased toward units with larger signals and higher firing rates, which are more likely to be deeper layer neurons. As discussed earlier, we observe the most inhibition in units in the deeper layers of PFC and MOs, where the other studies exclusively recorded, so perhaps this trend combined with the biased sampling of probes could explain the differences in results we observe.

Finally, these discrepancies could be explained by different effective strengths of CLA stimulation. The previous studies used viral injections, so their ChR2 expression likely reached higher levels compared to our transgenic Gnb4-IRES2-CreERT2;Ai32 mice. One possibility is that since our transgenic-based expression of ChR2 is likely weaker, the effective strength or number of spikes produced by our 5 ms optogenetic CLA stimulation is lower as well. In the CLA neurons we recorded from, the 5 ms optogenetic pulse triggered 1 spike on average in each opto-tagged CLA neuron, while in Jackson et al (2018), it appears that 2-3 spikes were triggered per CLA neuron. Perhaps if our ChR2 expression was stronger, light power was higher, or the duration of the pulse was longer, we could have triggered a similar blanket inhibitory state in the deep layers of PFC. In all, some combination of the above factors is likely to explain the differences we see between our study and previous work. However, the commonalities, including stronger input to cortical FS versus RS cells, likely reflect the robust aspects of the functional connectivity between CLA and cortex.

### Claustrum does not rhythmically entrain cortex at gamma frequencies

We find that 500 ms optogenetic stimulation of CLA causes the local circuit to oscillate in the gamma range (65-80Hz), which could emerge in part due to the strong connectivity between CLA interneurons (Kim et al., 2016), but this oscillation does not entrain downstream cortical neurons. We detect increased gamma power and phase synchronization between LFP signals in the downstream cortical areas, but no change in spike-LFP coupling, suggesting these are oscillations of presynaptic input from CLA axons that are coherent with each other because they originate from the same neural population (Schneider et al., 2021). We do not take this to mean that CLA definitely does not coordinate cortex; it is possible that since the mouse was not engaged in a task, the cortical areas were not primed for coordination. It is also possible that CLA does coordinate cortex, but at a different frequency or in an arrhythmic fashion.

Although our 20 Hz and 40 Hz optogenetic stimulations caused artifacts in the LFP band that precluded traditional coherence analyses, we did analyze pulse-wise spiking activity in response to these stimuli. We found that on average, the number of spikes following each pulse adapted more in response to 40 Hz pulses compared to 20 Hz pulses, suggesting lower frequencies may be more optimal for CLA-cortical activation. Furthermore, in the opto-tagged CLA neurons we recorded from, 20 Hz pulses on average produced facilitation of responses - the number of spikes tended to increase with successive pulses. This suggests that lower frequencies are more optimal for stimulating CLA neurons themselves, and perhaps a lower frequency than 20 Hz would have produced more facilitation in both CLA and downstream cortical neurons. This is consistent with previous work suggesting the CLA is involved in cortical slow-wave activity (Narikiyo et al., 2020).

In addition, even if CLA does not rhythmically coordinate cortex, our results show that it can coordinate average spike rates across different areas by causing a wave-like activation of neurons across the cortex (Supplemental Movies), as well as area-and layer-specific effects. This may reflect an important coordination whose function we do not yet understand.

### The function of claustrum

Our results show that simultaneously activating a significant fraction of CLA cells using optogenetics is not akin to a blanket inhibition of cortex as previously proposed (Jackson et al., 2018; Narikiyo et al., 2020). Indeed, our data clearly demonstrates that cortical RS cells, presumably largely cortical pyramidal neurons, can be excited by CLA activity.

Given the bi-directional connectivity between the CLA and almost all cortical regions (Atlan et al., 2016, 2018; Wang et al., 2017; Wang et al., 2022; Zingg et al., 2018), supplemented by widespread intra-claustral inhibition (Graf et al., 2020; Kim et al., 2016), it is possible that the CLA acts more like a hard or a soft winner-take-all (WTA) circuit (Grossberg, 1982; Itti et al., 1998; Maass, 2000), which could support the proposed role of CLA in salience processing (Remedios et al., 2014; Smith et al., 2019; Smythies et al., 2014). Consider the ecologically relevant situation in which the mouse must attend to one of several stimuli (possibly across different sensory modalities) or has to choose among one of several behavioral contingencies. Numerous populations of cortical neurons will be active under these conditions, relaying their information to the CLA where this activity competes, in an inhibitory manner, such that only the strongest activity survives within the CLA (or, that weaker cortical activity is severely attenuated in its CLA partner neurons). The output of the WTA is sent back to cortical targets, inhibiting all but one of the afferent cortical activity patterns; the one that is excited is the one that the animal attends (Atlan et al., 2018; Goll et al., 2015; Itti and Koch, 2001; White et al., 2020) and/or that the animal becomes conscious of (Crick and Koch, 2005). This model is consistent with findings that CLA perturbation affects behaviors that are more cognitively demanding (Atlan et al., 2018; Fodoulian et al., 2020; Terem et al., 2020; White et al., 2020), yet has little to no effect on simpler tasks (Remedios et al., 2010; White et al., 2020). Future experiments should be designed with these WTA dynamics in mind.

Our work emphasizes the importance of studying not just broad CLA-cortical connections, but connections between CLA and individual cortical areas and cell types, using, for instance, injection of retroAAV-Cre into specific cortical projection targets. It may be that CLA neurons perform distinct functions depending on the cortical area or areas to which they project. Evidence for this is emerging, such as in new work studying the roles of subpopulations of CLA neurons projecting to ACA versus ORB (Atlan et al., 2021), and to somatosensory cortex (Chevée et al., 2021). Our observation of the distinct functional impact of CLA on RSP, along with the lack of reciprocal connections from RSP to CLA, suggests this pathway may be particularly important to study as an output of the CLA. Even though the function or functions of CLA remain mysterious, its strong and widespread impact on cortical regions from the front to the back of the brain implies that it plays a vital role in cognitive and behavioral function.

## ACKNOWLEDGEMENTS

We thank the Allen Institute founder, Paul G. Allen, for his vision, encouragement, and support. We thank the Tiny Blue Dot Foundation for providing funding for this study. We thank the Allen Institute Laboratory Animal Services team for assistance with tissue collection, the Imaging team for assistance with imaging histology sections, and the Transgenic Colony Management team for generating and maintaining the mouse lines. We thank Quanxin Wang for useful feedback on the manuscript. We thank Sam Gale for use of his software for histology image processing. We thank Leslie Claar for useful discussions and assistance with setting up neurophysiology data processing.

## AUTHOR CONTRIBUTIONS

Conceptualization: E.G.M., D.R.O., C.K., and S.R.O. Methodology: E.G.M., J.R.K., D.R.O., A.A., and S.R.O. Investigation: E.G.M. and J.R.K. Data Curation: E.G.M. and J.R.K. Formal Analysis and Visualization: E.G.M. and S.R.G. Writing – Original Draft: E.G.M., S.R.G., and J.R.K. Writing – Review & Editing: all authors. Supervision: S.R.O., A.A., and C.K. Funding Acquisition: C.K.

## METHODS

### Mice

All mice were maintained by Allen Institute staff in the in-house animal facility and protocols were approved by the Allen Institute’s Institutional Animal Care and Use Committee. Thirteen Gnb4-IRES2-CreERT2;Ai32 mice, seven Esr2-IRES2-Cre;Ai32 mice, and two Cre-negative Gnb4-IRES2-CreERT2 mice (as controls) were used for this study. All mice were bred in-house. Gnb4-IRES2-CreERT2;Ai32 were induced with 5 days of tamoxifen at least 1 week prior to the initial implant surgery. After surgery, mice were single housed and maintained on a 12-hour reverse light cycle in a shared facility. Experiments were performed during the mouse’s dark cycle.

### Optic fiber cannula implant surgery

One to three hours before mice went into surgery, they were given the anti-inflammatory Dexamethasone (3-4 mg/kg IM) and the antibiotic Ceftriaxone (100-125 mg/kg SC). To anesthetize mice for surgery, they were initially placed in an induction chamber and induced with 5% isoflurane for approximately 1-3 minutes. Once anesthetized, the mice were placed in ear bars on a stereotaxic frame (Model 1900, Kopf) with an electric heating pad set to 37. 5°C to keep the animal’s body temperature constant. The mouse’s anesthesia was maintained with 1−2% isoflurane, and eye ointment (Systane Lubricant) was placed on the eyes to prevent them from drying out. Next, subcutaneous injections of either carprofen (5−10 mg/kg, SC) or ketoprofen (2-5 mg/kg, SC) were given to act as an analgesic along with atropine (0.02-0.05 mg/kg, SC) which decreased saliva production and helped regulate heart rate. Before an incision was made into the scalp, the surgical site was cleared of hair using scissors and Nair. The scalp was then cleaned three times, alternating between betadine and ethanol; the third and final coat of betadine was left on the scalp. Next, enough skin was removed from the scalp to accommodate the implant, the exposed skull was cleared of the periosteum, and the remaining skin was pushed to the sides of the skull. The mouse’s head was then leveled, the stereotaxic arm zeroed at bregma, and bregma was marked with a sharpie. Next, the stereotaxic arm was moved from bregma to coordinates x = − 3.7mm and y = +1.0 - 1.2mm; this location was marked with a sharpie, and the mouse’s head was tilted 10° to the right. A small hole was then drilled through the skull at the marked location with a medium drill bit (Carbide Friction 4 Grip 2mm 20.5in), and a blunt 27G needle was inserted 1.8 − 2.0mm down with the needle zeroed just above the brain.

After 10 minutes, the blunt needle was removed, and an optical cannula (Thorlabs Fiber Optic Cannula, Ø1.25 mm SS Ferrule, Ø400 μm Core, 0.39 NA, L=2 mm) was lowered 2.0mm into the brain. Next, a small piece of Gelfoam was placed in any gap between the skull and the cannula. The cannula was then secured to the skull with Metabond (Henry Schein MS) while ensuring the mark on bregma didn’t become obscured. The mouse’s head was then tilted back to its original center point. After re-zeroing the stereotaxic arm on bregma, a custom-made headpost was lowered onto the back of the animal’s skull. A custom 3D printed plastic well was then placed on the headpost, and both items were attached to the skull with Metabond. Finally, the mice were subcutaneously injected with up to 1ml of Lactated Ringer’s solution to replace any fluid lost during surgery and allowed to recover from anesthesia. Twice a day for two days after the mouse’s surgery, the animal received the same volume of injections as previously given of Carprofen and Ceftriaxone.

### Habituation to wheel and recording rig

One week after surgery, mice were habituated to being handled for 5 days by letting the mouse freely move in the habituator’s hands for five minutes each day. After each handling session, the mouse was given sunflower seed treats in their home cage. Once the mice became accustomed to handling, they were habituated to being head-fixed on a free-moving wheel. The amount of time the mouse spent head-fixed was gradually increased over 5 days from 5 to 30 minutes. The mice continued occasional 30-minute habituation sessions until the day before the experiment.

### Optogenetic perturbation and Neuropixels recording

On the day of or prior to the experiment, mice underwent craniotomy surgery to expose the brain for electrophysiological recording. As with the initial implant surgery, mice were given Dexamethasone and Ceftriaxone 1-3 hours prior, and then induced with 5% isoflurane for 1-3 minutes. Mice were then placed into the stereotax with an electric heating pad, anesthesia was maintained with 1−2% isoflurane, eye ointment was applied, and either carprofen or ketoprofen were injected subcutaneously along with atropine. After removing dental cement to expose the skull, bregma was located and used to guide the location of up to 3 craniotomies (<2mm diameter). After the brain was exposed, it was covered with kwik-cast for protection and to prevent drying between surgery and experiment. Mice recovered from surgery and anesthesia for at least 1 hour, and up to overnight, prior to placing them on the recording rig.

Prior to placing a mouse on the recording rig, probes were dipped in a fluorescent dye (CM-DiI, 1mM in ethanol, Thermo Fischer, V22888) to enable later locating them in histology. Next, mice were placed on the wheel and their headpost was fixed to an arm above the wheel. Running activity was recorded using a digital rotation encoder. The kwik-cast plug was removed, and agarose was placed into the well. The agarose mixture consisted of 0.4g high EEO Agarose (Sigma-Aldrich) 0.42g Certified Low-Melt Agarose (BioRad) and 20.5mL artificial cerebrospinal fluid (ACSF; 135mM NaCl, 5.4mM KCl, 1.0mM MgCl_2_, 1.8nM CaCl_2_, 5.0mM HEPES). A larger cone was lowered to protect the craniotomy and the probes, an optic fiber was attached to the optic fiber cannula, and a silver ground wire (32 AWG, A-M Systems) was placed into the agarose. Finally, additional ACSF was placed in the well to prevent the agarose from drying out.

Next, up to 3 Neuropixels probes (3a or 1.0) attached to individual 3-axis micromanipulators (New Scale Technologies) were lowered into place above the craniotomy. Probes were targeted to brain areas using bregma-to-motor coordinate transformations obtained by calibration. Once probes were located just above the brain surface, they were lowered steadily into the brain by NewScale software at a rate of 200μm/minute. Once probes reached their target depth (typically 2mm), they were allowed to settle for 10 minutes before initiating recording. Electrophysiological signals were recorded with OpenEphys software and were split into a spike band (500 Hz high-pass filter, 30 kHz sampling rate) and an LFP band (1 kHz low-pass filter, 2.5 kHz sampling rate). Acquisition occurred through either a dedicated FPGA board that streamed data over ethernet (Neuropixels 3a) or a PXIe card (Neuropixels 1.0).

The experiment proceeded as follows: First, a visual stimulus consisting of a flashing screen was presented to the mouse for 5 minutes. This screen was positioned to the right side of the mouse. Next, the optogenetic stimulation was initiated, with different patterns (5 ms, 20 Hz, 40 Hz, 500 ms) randomly interleaved with a 5 second inter-trial interval. At the end of the optogenetic stimulation (usually ~1 hour), the recording was stopped.

Most days, we performed 2-3 separate experiments on the same mouse by slowly retracting all probes and moving them to a new location, then repeating the steps above. In most cases, we recorded from mice on an additional subsequent day, for 2 total days of recording. At the conclusion of a day of recording, probes, ground wire, and optic fiber were removed, the craniotomy was re-sealed with kwik-cast, and the mouse was returned to its holding room. Probes were cleaned with 1% Tergazyme for at least 1 hour following the recording, then rinsed with DI water.

### Histology and Registration to Allen Common Coordinate Framework

After the end of all experiments, brains were removed in order to recover the locations of probes and the optic fiber implant. To prepare brains for histology, animals underwent cardiac perfusion with a 4% paraformaldehyde (PFA) solution. The extracted brains were placed in PFA for two days, rinsed with 1× PBS, and stored at 4°C in a foil-covered falcon tube filled with PBS.

Brains were sliced into 100μm sections using a vibratome (Leica VT1000 S), immediately mounted onto slides (Richard-Allan Scientific Slides), and dried on a slide holder with aluminum foil shielding them from light. After drying, the slides were coverslipped using Vectashield (Medium Mtg Vectashield Hardset DAPI) and sealed in the four corners with clear nail polish. The slides were then sent to the Allen Institute’s internal Imaging Core team to be digitally imaged. Slides were scanned with an Olympus VS110/120 fluorescent microscope at 10x magnification with DAPI (blue), FITC (green), and TRITC (red) filters.

The raw digital images received from the Imaging Core were processed according to the following steps. First, in custom python software, the images were cropped into individual sections, and their brightness was adjusted, so probe tracks were visible. Second, the individual slice images containing probe tracks were processed in Matlab using Sharptrack (doi:10.1101/447995). In Sharptrack, each slice was first aligned to the Allen Institute Common Coordinate Framework (CCFv3). Probe points were then visually identified and marked using the Sharptrack probe function. This step allowed estimation of the location in the CCF of each recorded unit. Because individual brains vary in size, as well as change in size due to fixation, the precise positions of the units were scaled based on two main criteria: 1) the relative low-frequency LFP power across channels, because a dramatic decrease in LFP power indicates the brain surface, and 2) anatomical landmarks such as ventricles or the corpus callosum where few to no units were expected. Finally, the resulting 3-d coordinate of each unit was entered as indices into the CCF reference volume to retrieve its estimated location in the mouse brain.

## Data processing

### Artifact masking & spike sorting

In many recordings, we observed an artifact associated with the optogenetic stimulation most likely due to the photoelectric effect. In the high-pass-filter spike band, these transient artifacts lasted approximately 1ms. These were more often observed with Neuropixels 3a versus Neuropixels 1.0 probes. In some cases, the amplitude of the artifact greatly exceeded that of most units. In such recordings, we masked the artifact by copying and reversing the voltage trace in the 1ms prior to the artifact. We then spike-sorted the data with Kilosort 2.0 and used quality metrics to determine good units (https://github.com/AllenInstitute/ecephys_spike_sorting). In recordings with a large artifact, we performed an additional manual curation of the Kilosort output to ensure no artifacts were counted as units.

### Local field potential processing

The low-pass filtered component of the signal, or local field potential (LFP) band, was more prone to optogenetic artifacts than the spike band. For most analyses, we limited the analysis window to 25-475ms after stimulation. Additionally, for both Neuropixels 3a and 1.0 probe recordings, we re-referenced the LFP signal to the median of the channels known to be outside of the brain, in ACSF. Unfortunately, due to the frequent turning on and off of the light during the 20 Hz and 40 Hz stimuli, we were not confident we could separate light artifacts from true signal.

## Data analysis

### Percent CLA illuminated

The CCF positions of implanted optic fiber cannula were estimated in the same way as probe tracks. With this information, we projected cones with different angles into the CCF volume in order to estimate the percent of CLA illuminated. With each cone projected into the volume, we computed the intersect between CLA and the light cone, then found the proportion of CLA voxels that were illuminated. 17 degrees was the angle obtained from using the diffraction index of the brain (1.34) and numerical aperture of the fiber (0.39). 26 degrees was a conservative estimate obtained from an empirical study of light spread using a fiber of identical diameter projected into a brain slice (Johnson et al., 2021). 45 degrees was a generous estimate obtained from figures of that same study.

### Running vs. stationary comparison

Mice were allowed to run or rest on a flat disc at will. Most mice rested on most trials (See Figure S2). When comparing running versus stationary trials, we observed a general decrease in the effect size of CLA stimulation on cortical neurons on running trials, likely due to the increase baseline firing rate during locomotion (Figure S2). Thus, unless otherwise noted, for all following analyses we used only stationary trials.

### Peri-stimulus time histograms and Z-scoring

We computed average peri-stimulus time histograms (PSTH) for each neuron and each condition by using the start times of each trial to extract spike times and then convert these to 1ms bin size histograms. Then, trials from a given condition were averaged together to produce the average PSTH. For each neuron, the mean and standard deviation of the spike counts across trials was computed, then used to produce a z-scored PSTH by first subtracting the baseline mean then dividing by the baseline standard deviation. In this way we could more easily compare activity of neurons with very different baseline firing rates and response sizes. For illustration, these normalized PSTHs were smoothed with a gaussian kernel.

### Fraction of neurons affected

To quantify the fraction of neurons affected in cortex by CLA stimulation, we computed the trial-by-trial spike counts in different windows, the duration of which was always matched between pre-stim and post-stim onset. Then for each neuron and condition, we performed a Wilcoxon sign-rank test on the trial-wise pre vs. post spike counts to obtain a p-value. Once all p-values were obtained across the dataset, we used the Benjamini-Hochberg correction for the false discovery rate with alpha set at 0.05. We then counted neurons with an adjusted p-value<0.05 as significantly affected, and determined whether they were excited or inhibited based on whether their spike count in the same window increased or decreased on average.

#### Time to Peak

To compute the time to peak for each neuron, we first smoothed the average PSTH with a gaussian kernel, then found the maximum of the PSTH within the first 75ms. We then used the estimated 3-d CCF coordinate for each neuron and the CCF coordinates of CLA to compute the Euclidean distance between each neuron and the nearest point of CLA. Finally, we plotted the time to peak versus distance to CLA for each neuron, broken down into various groups, and calculated the Pearson’s correlation and associated p-value across all neurons to determine whether time to peak correlates with distance from CLA.

#### Adaptation

To estimate the adaptation of neurons in response to the 20 Hz and 40 Hz pulse, we first found the spike counts in time windows 10-25ms following the times of each 20 Hz pulse, but across all conditions. In this way we could compare the same 10 spike counts across time and trials for each condition. To estimate the adaptation slope, we performed a linear regression for each neuron for each condition. We then could compare the distribution of slopes across neuron types and conditions.

### Functional clustering

#### Pre-processing

Neurons were clustered based on their mean response to 500 ms stimulation in two stages. For this, PSTHs at 1 ms resolution were first computed for all neurons and stimulus conditions, masking the time around stimulus onset and offset, and responsive units were identified based on criteria described earlier (Fraction of neurons affected). The PSTH for each neuron (including non-responsive ones) was convolved with a Gaussian window of width 4 ms and standard deviation of 2 ms and then z-scored with respect to 500ms of activity prior to the 500 ms stimuli.

#### Similarity clustering

The first 75 ms of responses were further down sampled by a factor of 3 to obtain response timeseries with 25 time points for each neuron. Cosine similarity was computed between all pairs of responsive neurons (n=3199). Each row of the similarity matrix was treated as a feature vector in n-dimensional space. The feature vectors were fit to a Gaussian Mixture Model with diagonal covariance and number of components ranging from 12 to 24 (*sklearn.mixture.GaussianMixture(n_components=<12-24>, covariance_type=‘diag’*)). Number of clusters was determined using the Bayesean Information Criterion (BIC), which ranged between 22 and 24 across repeated clustering attempts. The clusters were then merged based on an entropy criterion (Gouwens, 2020), leaving 22 clusters. Pairs of clusters with a similar distribution of pairwise similarities among constituent neurons within and across clusters (p>0.1) were further merged to obtain 19 clusters.

This process still resulted in a large number of clusters with very similar mean responses. Therefore, the above-mentioned clustering procedure was again applied to the mean responses of the clusters to finally obtain 9 clusters (Fig. S5A).

#### Functional connectivity

To estimate functional connectivity we used the FieldTrip package (Oostenveld et al., 2011) to compute the phase synchronization between the local field potential signal on pairs of channels. To reduce the impact of artifacts, but also include a large enough time window to examine phase synchronization at lower frequencies, we used a 25-475ms time window. In some experiments, the optogenetic-induced artifact was present even during this window. These recordings were excluded on the basis of computing the area under the curve of the LFP signals during optogenetic stimulation. Because the spatial scale of LFP signals is on the order of 100μm, we only included every fourth probe channel. To reduce the effect of volume conduction we used the debiased weighted phase lag index method (Vinck et al., 2011).

To examine whether spikes synchronized with oscillations in the local field, we used FieldTrip to compute the pairwise phase consistency (PPC) (Vinck et al., 2010). PPC was computed between each unit’s spike times and the average LFP from the nearest 4 channels that were at least 100μm away and within the same brain region. We then combined PPC spectra across units, and compared across units in different transgenic lines, cortical areas, and unit types. To investigate coupling between units, we used the MNE package in python (https://mne.tools) to compute spectral connectivity between pairs of neurons using binned spikes.

