## Supplemental Figures for "Influence of claustrum on cortex varies by area, layer, and cell type"

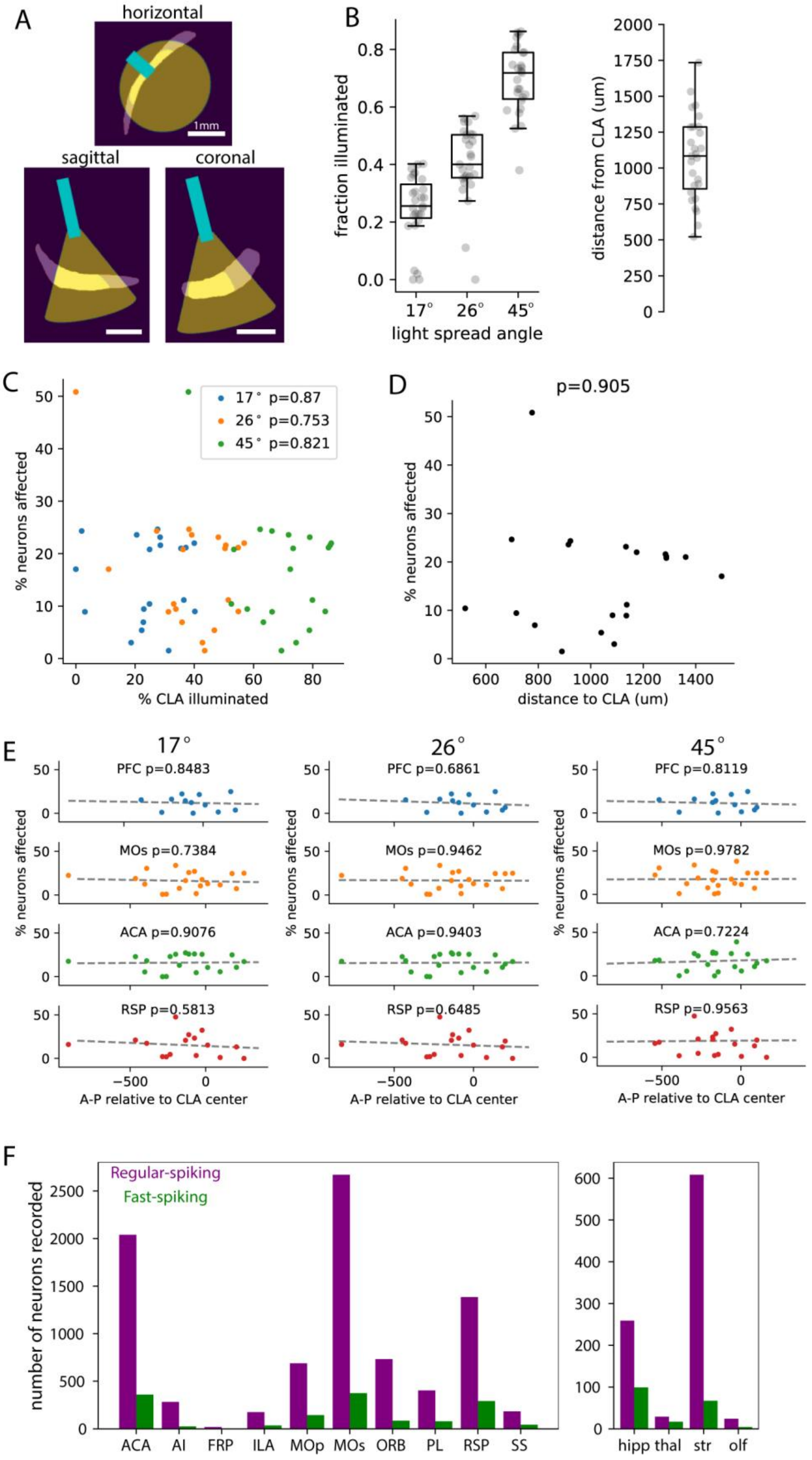

**Figure S1, related to Figure 1.**
**A**, example of a projected light cone based on the registered CCF position of an optic fiber implant in one experiment, shown in horizontal, sagittal, and coronal planes. The angle used for this projection was $26^{o}$. The CLA is shown in light purple, the optic fiber position in cyan, the projected light cone in yellow, and the estimated illuminated area of the CLA in bright yellow. Scale bar is 1 mm in CCF space. **B**, Left: estimated fraction of CLA illuminated across experiments from all mouse lines. Three different light spread angles were used, based on numerical aperture of the fiber and diffraction index of brain tissue ($17^{o}$), or published empirical measurements of light spread in brain tissue ($26^{o}$, $45^{o}$). Right: estimated distance of the tip of the optic fiber to the closest point of CLA. Boxplots show medians, quartiles, and 95% confidence intervals. **C**, scatter plot of percent neurons affected per experiment versus estimated percent of CLA illuminated. No significant trend was found (Pearson correlation). This could indicate that our higher estimates of the fraction of CLA illuminated are more likely to be correct, if the responses are close to saturation. The intensity of the illumination also likely decreases gradually, which would mean that changes in the angle and position of the cannula may not produce as much of a functional difference as the hard illumination cutoff of our simple estimation may suggest. **D**, scatter plot of percent neurons affected per experiment versus estimated distance of the optic fiber from CLA. No significant trend was found (Pearson correlation). **E,** scatter plots of percent affected neurons per experiment versus the center of the light stimulation in the A-P axis of CLA, separated by areas (rows) and the estimated angle of light spread (columns). No significant trends were found, suggesting differences in the A-P location of light stimulation in our experiments did not produce different spatial patterns of unit activation across different areas, despite the known topography of anterior CLA neurons projecting more to anterior cortex, and posterior CLA neurons projecting more to posterior cortex. **F**, left: number of units recorded across all cortical areas; right: number of units recorded in different categories of non-cortical areas.

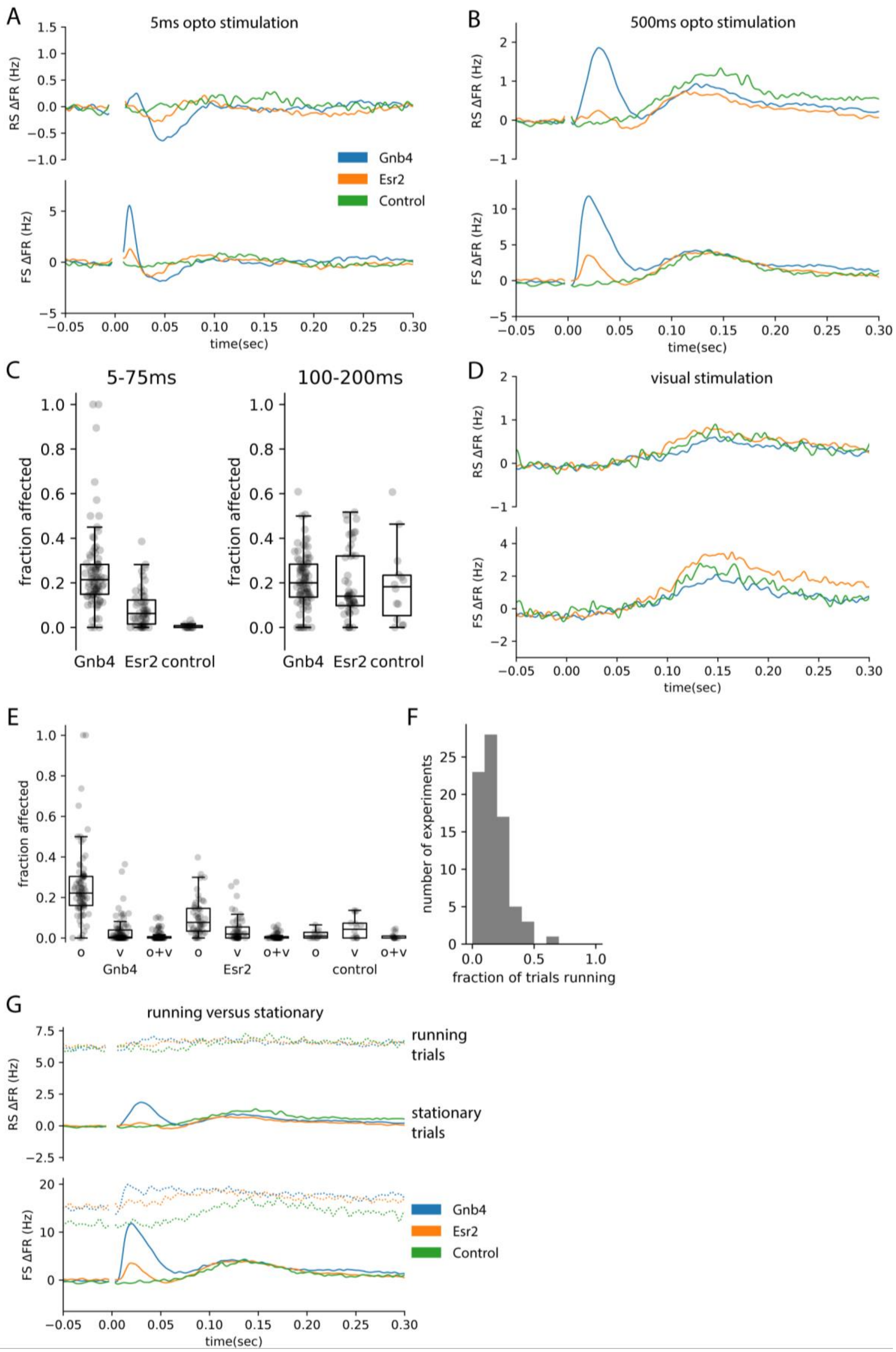


**Figure S2, related to Figure 2.**

**A**, average baseline-subtracted PSTHs of cortical units from the different transgenic lines (Gnb4, blue; Esr2, orange; control (no channelrhodopsin expression), green) in response to a 5 ms optogenetic stimulation. Upper, RS; lower, FS. **B**, same as A but for the 500 ms optogenetic stimulation. **C**, left: fraction of opto-affected units in the first 100 ms of 500 ms stimulation across all experiments separated by transgenic line; right: fraction of opto-affected units in the second 100 ms of 500 ms stimulation. **D**, average baseline-subtracted PSTH of responses of cortical units to a visual stimulus (full-screen grey to black, 250 ms duration). **E**, fraction of units affected only by opto stim (o), only by visual stim (v), or by both opto and visual stim (o+v). **F**, histogram of the fraction of trials spent running (>0.5cm/sec) across all experiments. **G**, same as A but also plotting average PSTHs during running trials (at least 5 running trials required to be included). Running PSTHs (dotted lines) from each neuron were normalized by subtracting the baseline firing rate during stationary trials, to show that the shift in baseline firing rate is as large or larger than the effect of CLA stimulation. Box plots show medians, quartiles, and 95% confidence intervals.

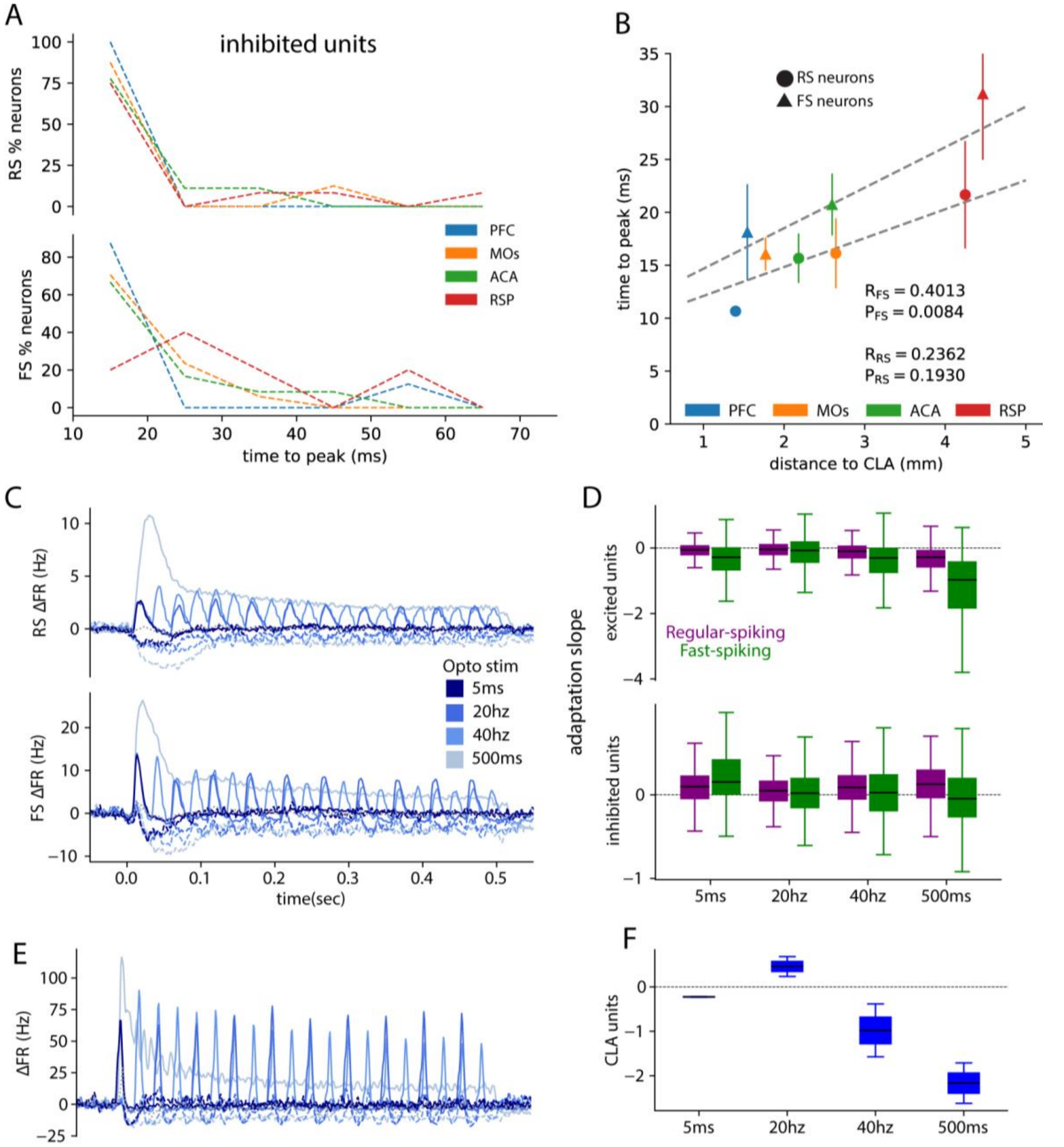


**Figure S3, related to Figure 3.**

**A**, histograms of time to peak firing rate for RS (upper) and FS (lower) neurons that were significantly inhibited in response to the 500 ms sustained stimulation, across the different cortical areas. **B**, Average time to peak for each area and cell type versus distance to CLA. The time to peak of FS units was significantly correlated with their distance from CLA (Pearson correlation). **C**, Average baseline-subtracted PSTHs for each optogenetic stimulation pattern across all RS (upper) or FS (lower) units. Units were split into categories according to whether they were significantly excited during the 500 ms sustained pulse (solid lines), inhibited (dashed lines), or not significantly changed (grey dotted line). **D**, boxplots of the adaptation slope of RS (purple) and FS (green) neurons across different conditions, separated by whether each neuron was initially excited (upper) or inhibited (lower). A negative slope indicates that a neuron's responses tend to decrease from the first to the last pulse. **E**, average PSTH of opto-tagged CLA units across the 4 stimulation patterns. **F**, boxplot of the adaptation slope putative CLA neurons across different conditions. Box plots show medians, quartiles, and 95% confidence intervals.

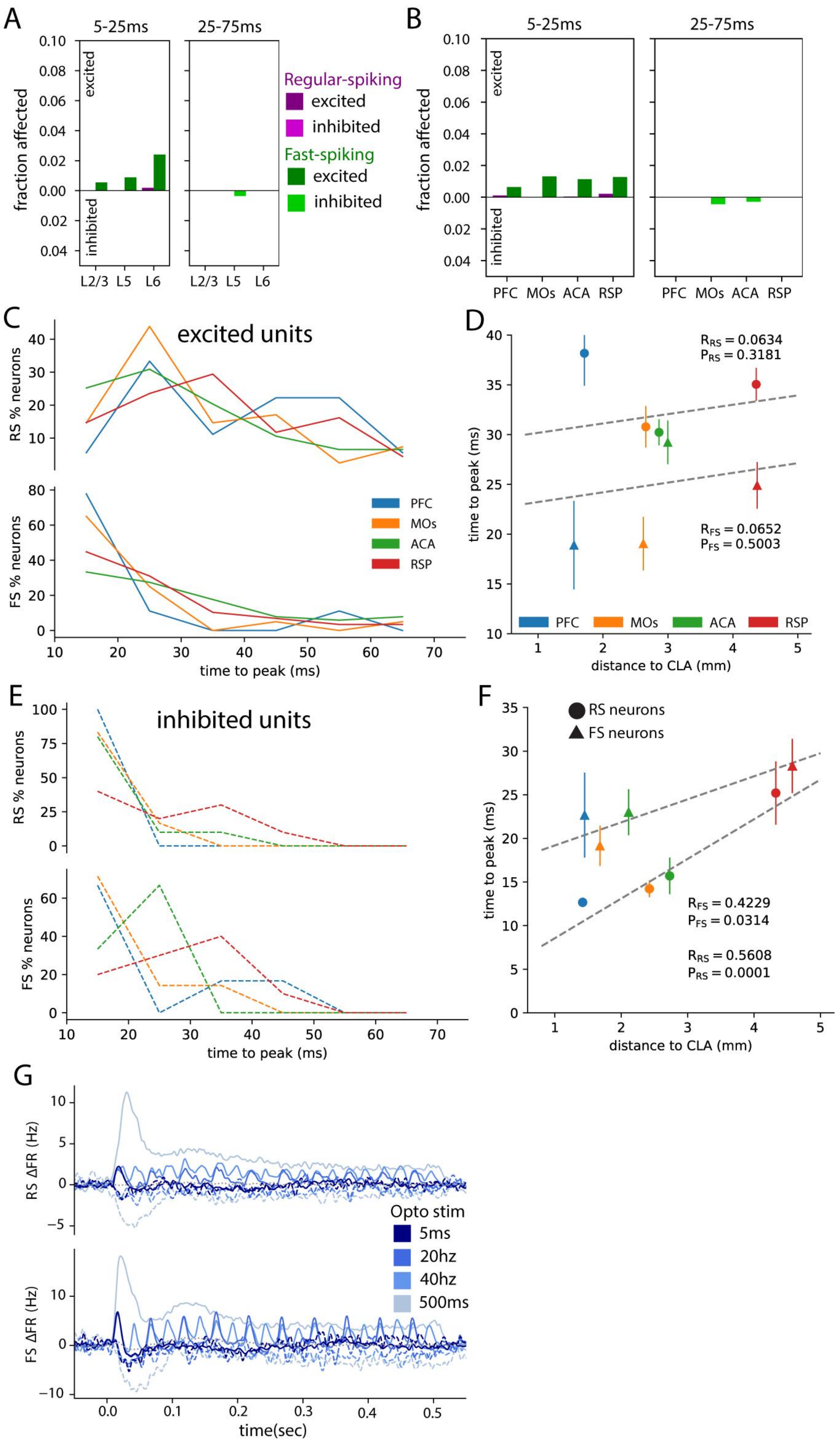


**Figure S4, related to Figure 4.**

**A**, fraction of neurons from Esr2-Ai32 mice significantly affected by the 5 ms perturbation within different time windows, separated by cortical layer. RS units are plotted in purple, FS units in green. The fraction of units with increased firing rates relative to baseline are plotted above 0, while units with decreased firing rates are plotted below 0. **B**, fraction of neurons from Esr2-Ai32 mice significantly affected by the 5 ms stimulation by cortical area. **C**, Time to peak histograms for significantly excited RS (upper) and FS (lower) neurons in response to the 500 ms sustained stimulation, across the different cortical areas. **D**, Average time to peak for each area and cell type versus distance to CLA. The time to peak of RS and FS units did not significantly correlate with their distance from CLA (Pearson correlation). **E,F**, same as C,D but for inhibited units. The time to peak of inhibited RS and FS cells did significantly correlate with distance from CLA. **G**, average baseline-subtracted PSTHs of units in response to the 4 optogenetic stimulation patterns across all RS (upper) or FS (lower) units. Units were split into categories according to whether they were significantly excited during the 500 ms sustained pulse (solid lines), inhibited (dashed lines), or not significantly changed (grey dotted line). Error bars SEM.

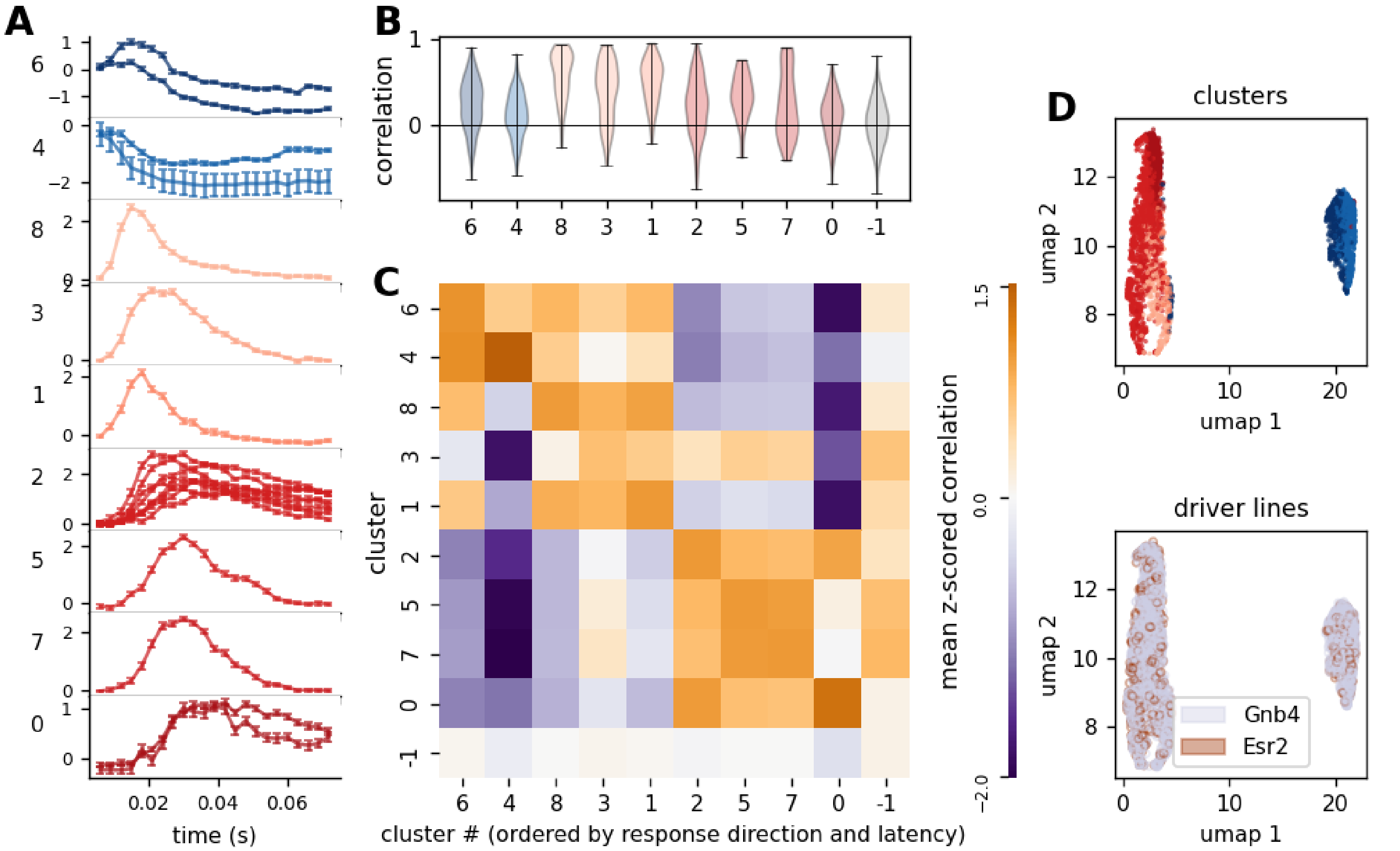


**Figure S5, related to Figure 5.**
Details of clustering. **A,** Mean responses of the 19 preliminary clusters of response types. These were further clustered based on similarity to obtain 9 distinct response types (each panel corresponds to one of the 9 final response types, showing the mean responses of the 19 original clusters that were grouped together; cluster number indicated to the left). **B,** Distribution of correlations between response of each neuron to the mean response of the parent cluster for the 5 ms stimulus. Although clustering was performed on the 500 ms responses, the 5 ms responses also show high fidelity to the clusters. **C,** Average correlation between responses (to 500 ms stimulation) of individual neurons and cluster mean responses. Responses of neurons in clusters (4, 6), (8, 3, 1) and (2, 5, 7) are similar to one-another but well-separated from other clusters. **D,** UMAP performed directly on response timeseries (as opposed to UMAP performed on the feature vectors, as shown in Figure 5) shows same large clusters (inhibited and excited) as UMAP on feature vectors (Figure 5A,C). Note the intermixing of Gnb4 and Esr2 cells in both large clusters.

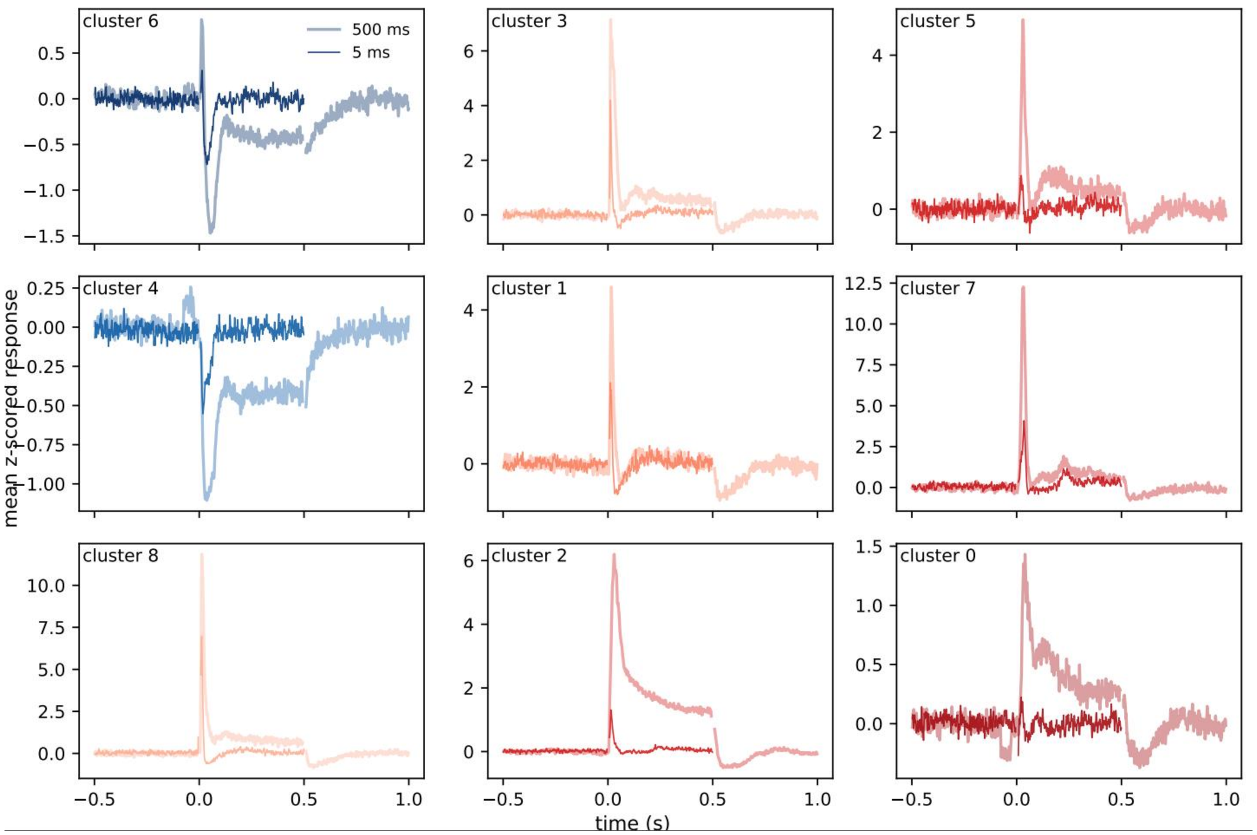


**Figure S6, related to Figure 5.**

Similarity between mean 500 ms and 5 ms responses of each cluster. Clusters 8, 1 and 3 show a clear inhibition followed by the initial excitation in the 5 ms response. The inhibition tends to be overridden by the continuous drive in the 500 ms stimulation for clusters 8 and 9, but is still visible in cluster 1. Clusters 2, 7 and 0 do not have any inhibition.

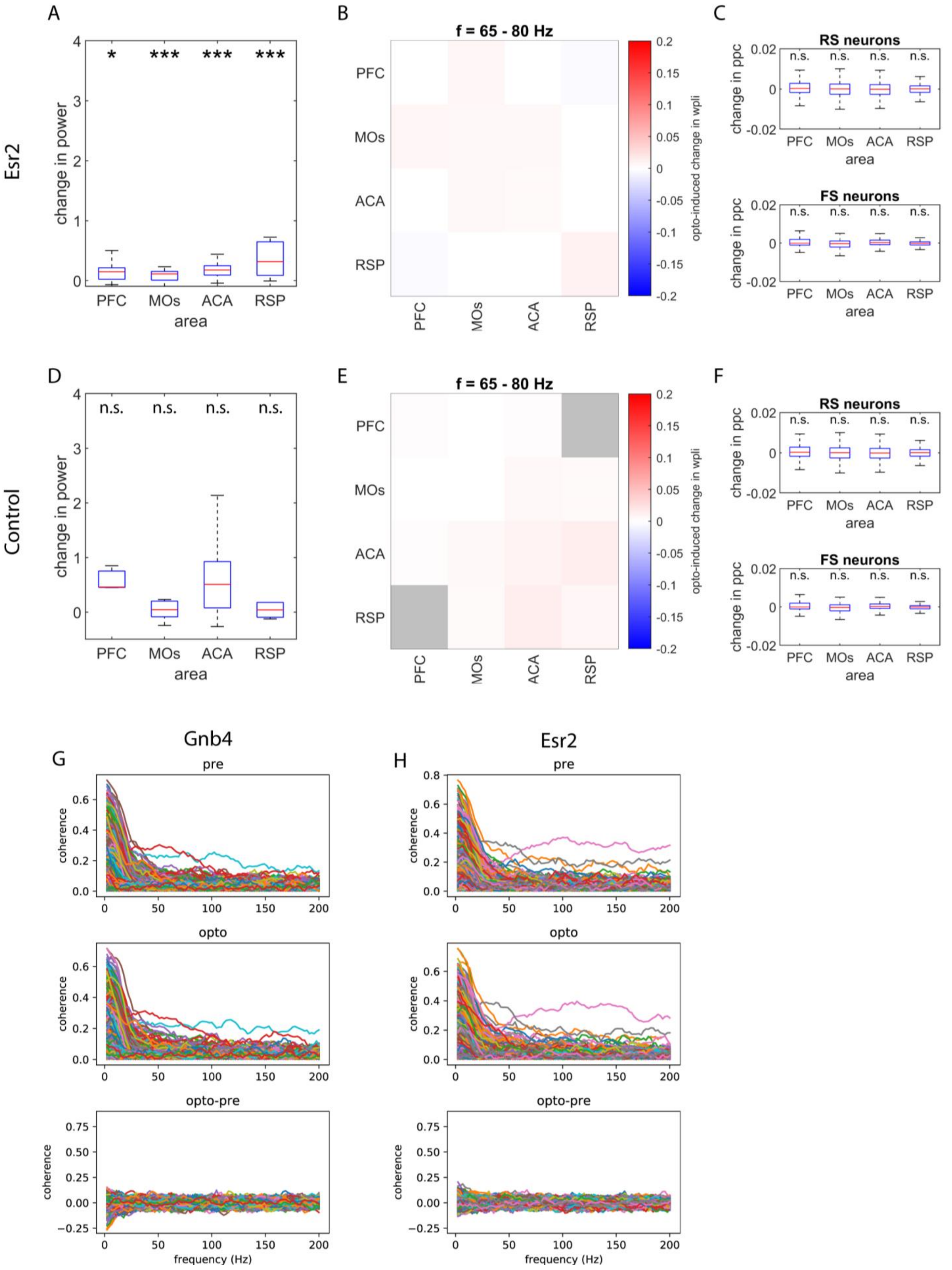
A
B
C
Esr2
change in power
* *** *** ***
PFC MOs ACA RSP
area
f = 65 - 80 Hz
opto-induced change in wpli
RS neurons
FS neurons
change in ppc
n.s. n.s. n.s. n.s.
D
E
F
Control
n.s. n.s. n.s. n.s.
G
H
Gnb4
Esr2
pre
opto
opto-pre
coherence
frequency (Hz)

**Figure S7, related to Figure 6.**
**A,** summary of CLA-stimulation-induced change in power in Esr2 mice, across different areas. Significant changes in the gamma band (65-80Hz) were found in all areas. **B,** heatmap of changes in spectral connectivity between pairs of areas in Esr2 mice. We did not find significant changes in the 65-80hz band. **C**, summary of changes in pairwise phase consistency across areas and cell types in Esr2 mice. We did not observe any significant changes in this measure. **D-F,** same as **A-C**, but for control mice. **G**, coherence spectra among pairs of cortical neurons from Gnb4 mice prior to (pre) and during optogenetic stimulation of CLA (opto), and the difference between them (opto-pre). This demonstrates that even though many pairs of neurons display low-frequency coherence with each other, our 500 ms CLA stimulation did not alter this coherence. **H**, same as G but for unit pairs recorded in Esr2 mice. Box plots show medians, quartiles, and 95% confidence intervals.

**Supplemental Movie 1: RS units from Gnb4-Ai32 mice**
Cortical RS neurons recorded in Gnb4-Ai32 mice that were significantly excited or inhibited by 500 ms claustrum stimulation (see Methods) are projected onto a sagittal view of the mouse brain atlas, preserving their relative anterior-posterior and dorsal-ventral positions. CLA is highlighted in light gray. A small amount of random jitter was added to the A-P coordinate of each neuron to increase visibility of individual points. Each axis increment corresponds to 10 microns, as the 10-micron resolution atlas was used for probe and unit location registration. Each point corresponds to one neuron. As the movie plays, the color of each neuron changes, corresponding to that unit’s average normalized firing rate in response to 500 ms claustrum stimulation, with red representing an increase and cyan a decrease in activity. Note, when the optogenetic stimulus activates at time=0, a slight onset artifact may be observed. As noted elsewhere, the 5 ms before and after the onset of the optogenetic stimulus were excluded from quantitative analyses.

**Supplemental Movie 2: FS units from Gnb4-Ai32 mice**
Same as Movie 1 but plotting FS neurons that were either significantly excited or inhibited by 500 ms claustrum stimulation.

**Supplemental Movie 3: RS units from Esr2-Ai32 mice**
Same as Movie 1 but plotting neurons recorded in Esr2-Ai32 mice.

**Supplemental Movie 4: FS units from Esr2-Ai32 mice**
Same as Movie 2 but plotting neurons recorded in Esr2-Ai32 mice.
